# Parasites make hosts more profitable but less available to predators

**DOI:** 10.1101/2022.02.08.479552

**Authors:** Loïc Prosnier, Nicolas Loeuille, Florence D. Hulot, David Renault, Christophe Piscart, Baptiste Bicocchi, Muriel Deparis, Matthieu Lam, Vincent Médoc

## Abstract

Parasites are omnipresent, and their eco-evolutionary significance has aroused much interest from scientists. Parasites may affect their hosts in many ways with changes in density, appearance, behaviour and energy content, likely to modify their value to predators (profitability) within the optimal foraging framework. Consequently, parasites could impact predators’ diet and the trophic links through food webs. Here, we investigate the consequences of the infection by the iridovirus Daphnia iridescent virus 1 (DIV-1) on the reproductive success, mortality, appearance, mobility, and biochemical composition of water fleas (*Daphnia magna*), a widespread freshwater crustacean. We do predation tests and compare search time, handling time and feeding preference between infected and uninfected *Daphnia* when preyed upon by *Notonecta* sp., a common aquatic insect. Our findings show that infection does not change fecundity but reduces lifespan and thereby constrains fitness. Infected *Daphnia* show reduced mobility and increased color reflectance in the UV and visible domains, which potentially affects their appearance and thus vulnerability to predators. Infection increases body size and the amount of proteins but does not affect carbohydrate and lipid contents. Although infected *Daphnia* are longer to handle, they are preferred over uninfected individuals by aquatic insects. Taken together, our findings show that DIV-1 infection could make *Daphnia* more profitable to predators (24% energy increase), a positive effect that should be balanced with a lower availability due to the higher mortality of infected specimens. We also highlight that exposure to infection in asymptomatic individuals leads to ecological characteristics that differ from both healthy and symptomatic infected individuals.

**Recommender:** Luis Schiesari

**Reviewers:** Thierry De Meeus and Eglantine Mathieu-Bégné

**Cite as:** Prosnier L., N. Loeuille, F.D. Hulot, D. Renault, C. Piscart, B. Bicocchi, M, Deparis, M. Lam, & V. Médoc. (2023).

Parasites make hosts more profitable but less available to predators. BioRxiv, ver. 4 peer-reviewed and recommended by Peer Community in Ecology. https://doi.org/10.1101/2022.02.08.479552

## Introduction

All living organisms are concerned by parasitism, either as hosts or because they practice this strategy themselves at some point in their lifecycle (Dobson et al., 2008). Infection is generally accompanied by subtle or severe alterations in host phenotypes, including changes to physiology, morphology, and behavior with potential consequences on fitness (Thomas et al., 2010). Host fitness can be impacted directly through reduced fecundity or increased mortality, or indirectly when phenotypic alterations make the hosts more vulnerable to their natural enemies, including predators. Only few studies working on the diversity of parasite-induced phenotypic alterations have simultaneously considered both direct and indirect effects (see the review of Cézilly et al., 2013). From the predators’ perspective, their fitness can also be indirectly reduced by infection of their prey, leading to the possible avoidance of infected prey (see the meta-analysis of Flick et al., 2016).

The direct effects of infection result from the rerouting of metabolic energy from the host to parasite growth, maturity, and reproduction, with the intensity depending on parasite virulence. Virulence can be defined as the extent to which a parasite exploits its host and thus reduces its survival and fecundity (Read, 1994). Owing to its importance, virulence is very often assessed in host-parasite interactions (Prins & Weyerhaeuser, 1987; Newey & Thirgood, 2004). For instance, some parasites of water fleas (e.g., fungus, bacteria, trematode) reduce egg production and increase mortality (Schwartz & Cameron, 1993; Decaestecker et al., 2003). Host survival can also decrease indirectly (i.e., implying a third species) when infected hosts become less competitive (Decaestecker et al., 2015), or more vulnerable to predation – which is either considered adaptive from the point of view of the parasite when the predator is the next host (see the manipulation hypothesis, Bethel & Holmes, 1977; Lefèvre et al., 2009; Jacquin et al., 2014), or a simple by-product of infection. For instance, the reduced body condition of infected moose makes them more prone to be eaten by wolves (Peterson & Page, 1988), while infected red goose are more readily attacked by mammalian predators (Hudson et al., 1992). Similarly, infection with the nematode *Gasteromermis* sp. reduces larval drift in the insect *Baetis bicaudatus*, which becomes more vulnerable to predation by the sickle springfly *Kogotus modestus* but not to predation by the caddisfly *Rhyacophila hyalinata*, thus suggesting a predator-dependent effect (Vance & Peckarsky, 1997). Host weakening (see the review of Sánchez et al., 2018) may be due to energy reallocation to parasite growth (Hall et al., 2007) or to the cost of the immune response (Otti et al., 2012). Increased vulnerability can also result from changes in host appearance (e.g., coloration, size). For instance, *Polycaryum laeve* (Chytridiomycota) infection causes opacification in *Daphnia pulicaria*, which may increase its vulnerability to fish predation (Johnson et al., 2006).

Parasite-induced phenotypic alterations in prey are likely to influence the diet of predators. Optimal foraging theory predicts that the inclusion of a particular prey to the diet of a predator depends on its relative abundance and profitability ranking (Emlen, 1966; MacArthur & Pianka, 1966; Charnov, 1976a; b). Profitability is the ratio between the energy content of the prey and its handling time for a given search time. By diverting energy, parasites modify the biochemical content of their host. In particular, Plaistow et al. (2001) reported a decrease in glycogen content and an increase in lipid content in crustacean amphipods infected by the acanthocephalan parasite *Pomphorhynchus laevis*. For *Daphnia pulicaria* infected by *Polycaryum laeve*, the increase in carbon content and the reduction in nitrogen and phosphorus increased the carbon-to-nitrogen ratio (Forshay et al., 2008). When energy content is increased by infection, hosts might conversely become more profitable to predators if the handling time remains unchanged. Similar effects are expected when alterations in behavior and aspect make host weaker (reducing prey escape) and more visible, and thus more vulnerable (lower search time and handling time) to predation.

To understand the effects of parasitism in a trophic context, it is crucial to study concomitantly the different host alterations and their relative intensity. To address this issue, we used as host species the water flea *Daphnia magna*, a widespread freshwater crustacean that plays a central role in food webs, both as an herbivore and as a prey (Lampert & Sommer, 2007; Reynolds, 2011; Ebert, 2022). *Daphnia magna* can host a diversity of parasites (Green, 1974; Ebert, 2005, 2022), including the Daphnia iridescent virus 1 (DIV-1, Toenshoff et al., 2018), which is known to increase mortality, reduce fecundity (Ebert et al., 2000) and alter activity, potentially affecting their profitability to the predators that do not risk infection by this highly specific parasite. DIV-1 also impacts host appearance through the induction of a white phenotype and, consequently, has been known as “White Fat Cell Disease” (WFCD) but wrongly labeled as “White Bacterial Disease” (WBD). However, information on the phenotypic modifications and their implications regarding vulnerability to predation are lacking, which prevents us from fully understanding the consequences of parasitism in an optimal foraging context. We quantified the alterations in terms of fecundity, survival, mobility, coloration, body size, biochemical content (carbohydrates, lipids, and proteins), and vulnerability to predation (by *Notonecta*, a common generalist predator (Giller, 1986; Van der Lee et al., 2021) and fish) using both *in situ* and experimentally-infected *D. magna*. Considering previous research on the virulence of DIV-1 (Ebert et al., 2000), we expect strong direct effects with a reduction in host survival and fecundity. Indirect effects are studied here for the first time, and we expect the energy costs of infection to reduce host activity, thus favoring predation, which could be further facilitated by the white coloration of infected water fleas.

## Material and Methods

### Collection and maintenance of organisms

*Daphnia magna* (identified according to the morphological characteristics described by Amoros, 1984) and the parasite were collected from two ponds in Paris (France): La Villette (48°53’43.0“N 2°23’26.5“E) and Bercy (48°50’03.0“N 2°23’03.1“E) where DIV-1 prevalence ranges from 0.5 to 3% (pers. obs.). Given the high host specificity of DIV-1, collecting hosts and parasites from the same pond was expected to promote the success of the experimental infection (Decaestecker et al., 2003). DIV-1-infected *D. magna* have a highly identifiable phenotype (Fig. C1): under light, infected fat cells are blue-white, almost fluorescent (Ebert, 2005). This white phenotype is highly characteristic to an iridovirus, and only one, the DIV-1, was recently identified by Toenshoff et al. (2018). They used only one Finland population for the determination but found that this highly specific parasite also infects *D. magna* from European ponds (e.g., in France), known to have individuals showing the White Fat Cells Disease. Thus, it is likely that our specimens displaying the White Fat Cell Disease (i.e., the white coloration) were infected with DIV-1.

All *D. magna* individuals were stored in 5-L rearing tanks (100-150 ind.L^-1^) filled with filtered water from their collection pond. Depending on the experiment, they were used on the day of capture or stored for up to 3 days without food supply at 20 °C. To identify infected individuals and isolate parasites, the crustaceans were placed in a black jar and illuminated to observe any phenotypic signs of infection. Infected and non-infected *D. magna* were kept separately in Volvic® mineral water at 20°C under a 12:12 light:dark cycle (200 Lux) at the same density of 100 ind.L^-1^ in 1-L tanks.

Vulnerability to predation was investigated using an aquatic insect from the *Notonecta* genus and a fish, the European minnow *Phoxinus phoxinus* (Appendix A). *Notonecta* sp. (1.8-2.0 cm in total length) were collected from a pond at Orsay (France, 48°42’04.4“N 2°10’42.7“E) using a hand net. Immediately after collection, they were stored and starved in 5 L of water from the pond (3 ind.L-1) for 1 day before the beginning of the experiments.

In this study, we performed an experimental infection to determine the effects of DIV-1 on fecundity (Measure 1), mortality (Measure 2), mobility (Measure 3), and size (Measure 4). We also used naturally-infected individuals to measure fecundity (Measure 1), mobility (Measure 3), size (Measure 4), energy content (Measure 5), coloration (Measure 6), vulnerability to predation (Measure 7&8), and predator preference (Measure 9). Table 1 and Fig. C2 summarizes the measures performed on each collected Daphnia.

**Table 1.**
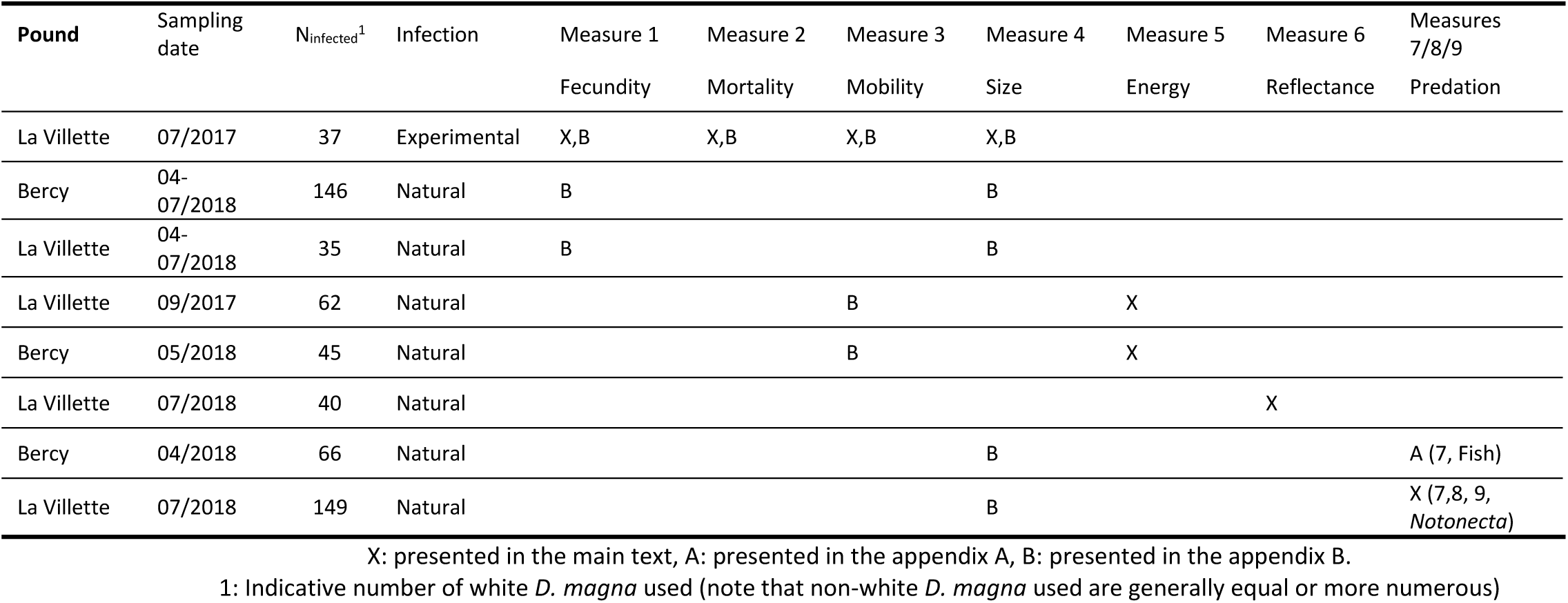
Summary of measurements performed for each collected *D. magna*.

### Fecundity and mortality (Measures 1 and 2)

Reproductive success (Measure 1) and survival (Measure 2) were assessed in two manners: in the laboratory through experimental infections (Measures 1 and 2) and from wild individuals (Measure 1). The experimental infection allowed us to clearly distinguish between the effects on fecundity and survival. We do not consider offspring production along lifetime as a proxy of fecundity, but rather as a proxy of fitness, because it encapsulates both fecundity parameters (clutch size, clutch frequency, and age at maturity) and survival (lifespan).

Gravid *D. magna* collected from the La Villette pond in July 2017 and stored in their rearing tanks were transferred individually to 50-mL jars containing Volvic® water. Newborns (<24h) were transferred individually into jars with 45 mL of Volvic® water in a climatic chamber at 20 °C, and fed with 0.25 mL of *Scenedesmus obliquus* (2.3×10^6^ cells.mL^-1^) every 3 days throughout the experiment. These algae were obtained from the Muséum National d’Histoire Naturelle (Paris, France, algothèque MNHN; strain number: ALCP n°349), and cultivated at 20 °C under a 12:12 light:dark cycle in an ES medium (Basal Medium, “Erddekokt + Salze“ described by Culture Collection of Algae of Sammlung von Algenkulturen Göttingen). Molts were removed daily to maintain water clarity.

To infect *D. magna*, we prepared a solution of infected *D. magna* cadavers (hereafter, parasite solution) homogenized at the concentration of 1 cadaver/mL in Volvic® water. We used individuals infected naturally and showing the white phenotype. A control solution was prepared with healthy cadavers (i.e., with individuals not showing the white phenotype). Half of the newborns were exposed to the parasite solution and the other to the control solution. On Day 1, we added 1 mL of the solution to obtain a ratio of 1 cadaver *per* juvenile of *D. magna*. On Days 4 to 6, we stirred the water (both the control and treatment) using a pipette to resuspend the spores and promote infection. Water was replaced on Day 15 by clean water (without the virus) and then once a week until the death of the last individual of *D. magna* (163 days). Offspring were removed and counted daily, and dead *D. magna* were controlled visually, as described above, for infection signs. We started two sets of experimental infections with 1 day of delay: the first set was performed with 27 juveniles (14 exposed to the parasite solution and 13 to the control solution) coming from 11 distinct mothers, while the second set was performed with 44 juveniles (23 exposed to the parasite solution and 21 to the control solution), also coming from 11 distinct mothers. The experiment lasted until the death of all *D. magna*, representing 163 days. We also measured the fecundity of naturally-infected individuals (see Appendix B).

### Mobility (Measure 3)

We assessed mobility in two ways: (i) using the experimentally-exposed individuals from Measure 1 that were still alive on day 14 (n = 53), and (ii) using naturally-exposed individuals (see Appendix B). These naturally-infected individuals were subsequently used for Measure 5 (see below). We measured speed (maximum and mean), swimming time, and the number of turnings as described by Untersteiner et al. (2003) and Bownik (2017). The water fleas were placed individually into one of the nine chambers (3 x 3.2 x 1 cm, L x l x h) of a grid in a black box filled with Volvic® water. We placed a light source (150 Lux) under the grid with a video camera (Canon® EOS 70D body with Canon® EF-S 17-55mm f/2.8 IS USM lens) placed 52 cm above. After 5 min of acclimatization, *D. magna* were filmed for 29 sec, divided into five sequences of 3.80 sec, each interrupted by 5 sec intervals between two consecutive sequences, in monochrome at a rate of 25 fps. By making five films *per* animal, we reduced the risk of misdetection by the software. Several sequences in which *D. magna* were not detected were not analyzed, and mobility was instead evaluated in the three or four remaining films. Video analysis was performed with the ImageJ software (version 1.4.3.67) and the plugin wrMTrck (31/10/2011 version by Jesper Søndergaard Pedersen, modified by the authors). We subtracted the background and shifted from grayscale to black and white to promote detection. The plugin allowed us to identify the group of black pixels corresponding to *D. magna* and determine the mobility parameters (mean and maximum speeds, rotating movements). We modified the plugin to assess inactivity time: the absence of movement between two consecutive records was converted in time by considering the time interval between these two sequences (here 1/25 sec).

### Body size (Measure 4)

To measure individual size (from the head to the start of the caudal spine) of the experimentally-infected *D. magna* used for Measures 1 & 2, we used the video recordings obtained for the assessment of mobility (Measure 3, n = 53 individuals) (see Appendix B for naturally-infected individuals). We also used the photographs of a set of *D. magna* used in the predation experiments (Measure 7, see below, n = 229) to determine their size. Specimens of *D. magna* taken from photographs and videos were measured with ImageJ software (version 1.4.3.67).

### Biochemical composition and energy value (Measure 5)

We assessed the quantity of carbohydrates, lipids, and proteins *per* mg of *D. magna* in the naturally-infected *D. magna* used for Measure 3. For each pond, we considered three categories of crustaceans: broodless individuals (no visible signs of infection, no eggs), brooding individuals (no visible signs of infection, with eggs), and infected individuals (visible signs of DIV-1 infection with the white coloration, without eggs). Unfortunately, we did not collect enough DIV-1-infected *D. magna* with eggs to conduct biochemical assays. Preliminary tests showed that pools of 10 individuals were optimal to obtain a reliable signal for accurately measuring the amount of proteins, sugars, and triglycerides. Immediately after the mobility experiment, groups of 10 *D. magna* individuals were snap-frozen and stored at -25 °C after removing water with a towel.

The concentrations of proteins, sugars, and triglycerides were measured using colorimetric assays, as described by Ouisse et al. (2017) and Foray et al. (2012). Briefly, each pool of 10 crustaceans was first weighed (Fresh mass, Balance XP2U Mettler Toledo, Columbus, OH, d=0.1 µg). After the addition of 200 µL of phosphate buffer (pH 7.2), each pool was homogenized for 90 sec at 25 Hz (bead-beating device, Retsch™ MM301, Retsch GbmH, Haan, Germany). The pools were then centrifuged (180 g, for 10 min, 4 °C), and a volume of 8 µL of supernatant was collected to quantify the amount of proteins using the Bradford method (Bradford, 1976). The absorbance of samples was read at 595 nm, and the protein concentration was calculated from the calibration curve from different concentrations of bovine serum albumin.

The rest of the supernatant (192 µL) was mixed with 148 µL of phosphate buffer and 510 µL of a methanol-chloroform solution (ratio 2/1, volume/volume). After centrifugation at 180 g and 4 °C for 10 min, 15 µL of chloroform was transferred to the new microtubes for the triglyceride assays and stored at -20 °C. The pools were redissolved into 200 µL of Triton-BSA buffer. The manufacturer’s instructions were followed for the triglyceride colorimetric assay (Triglycerides, kit reference CC02200, LTA SRL, Italy).

For the measurement of total sugars, 80 µL of the methanol-chloroform solution of each pool were dried for 30 min at room temperature before adding 300 µL of fresh anthrone solution (1.42 g.L^-1^ anthrone in 70% acid sulfuric solution). Next, the pools were heated at 90 °C for 15 min, and the absorbance was measured at 625 nm. Different glucose concentrations were used for drawing the calibration curve, and total sugar amounts were thus expressed as glucose equivalents.

We then calculated total energy content, in mJ, using the energy of combustion (Gnaiger, 1983; de Coen & Janssen, 1997): 17,500 mJ.mg^-1^ glycogen, 39,500 mJ.mg^-1^ lipid, and 24,000 mJ.mg^-1^ protein. We summed the three energy contents to determine the energy, in mJ, *per D. magna* and *per* mg of *D. magna* (i.e., taking into account the mass differences between each type of individuals).

### Reflectance (Measure 6)

We measured *D. magna* reflectance around the midgut where the parasite-induced alteration in coloration is observable using a spectrophotometer (USB2000+) between 280 and 850 nm (DH-2000 Deuterium Tungsten Source, 210-1700nm), and the SpectraSuite Cross-Platform Spectroscopy Operating Software. We used 80 naturally-exposed *D. magna* (40 presenting no visible sign of infection and 40 with a visible white coloration) collected in July 2018 from the La Villette pond and kept in rearing tanks for less than 6 hours. We alternately measured five uninfected and five infected *D. magna,* removing water with a towel for a few seconds before the measurement.

### Susceptibility to insect predation (Measures 7 and 8)

*Notonecta* sp. (n = 13) were starved for 24 h before the experiments, and *D. magna* were collected from the La Villette pond in July 2018 and used within 6 hours. We used 500-mL jars filled with spring water (Cristaline®, Cristal-Roc source) and performed a first experiment on the timing of capture and handling time (Measure 7 & 8) and a second experiment on prey choice (Measure 9).

For the timing of capture (Measure 7), after 24 h of acclimatization for the *Notonecta* sp., we introduced three *D. magna* either infected or presenting no sign of infection (hereafter healthy). During 1 hour, we recorded the times of capture of alive prey and the release of each prey cadaver. We defined handling time (Measure 8) as the time interval between capture and release, and intercapture time as the time interval between the release of the current prey (or the start of the experiment) and the capture of the next prey. We simultaneously offered healthy *D. magna* to half of the *Notonecta* sp. and infected *D. magna* to the other half. After another 24 h period of acclimatization and starvation, we performed the same experiments with the other prey type *per* predator.

To investigate prey choice (Measure 9), we offered 10 healthy and 10 infected *D. magna* to each of the 13 *Notonecta* sp. after a 24-h period of acclimatization and starvation. When approximately half of the prey was consumed, we stopped the experiment, counted the surviving *D. magna*, and identified their infection status. To determine the preference of the predator for infected prey, we used the Manly’s alpha index (Manly, 1974; Goren & Ben-Ami, 2017).

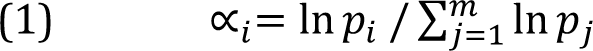

where *∝_𝑖_* is the Manly’s alpha for prey type *i* (the infected prey here), 𝑝_𝑖_ and 𝑝_𝑗_ are the proportions of prey types *i* and *j*, respectively, at the end of the trial, and 𝑚 is the total number of prey type (here 2). If *Notonecta* sp. prefers infected *D. magna*, then *∝_𝑖_* tends to 1, a *∝_𝑖_* value of 0.5 indicating the absence of preference.

### Statistical analyses

Statistical analyses were performed using R (version 3.4.3) with a significance threshold of 5%, and summarized in Fig. C2. We used a Multiple Factor Analysis (MFA) to analyze the fecundity, survival, size and mobility (Measures 1-4) of experimentally-infected daphnia with 10 parameters aggregated in four factors: Clutch Size/Clutch Frequency/Maturity (Fecundity), Lifespan (Lifespan), Maximal Speed/Average Speed/Number of Turns/Inactivity (Mobility), and Size (Size). Because total egg production results from a combination of fecundity and lifespan traits, it was added as a supplementary parameter as well as the status of infection. In addition to the MFA, we also analyzed separately these 10 parameters to compare with the results obtained with naturally-infected individuals (see Appendix B).

The biochemical composition (Measure 5) was analyzed using ANOVA and two-sided pairwise t-tests of Welch using the Holm adjustment method because the residuals were normally distributed, sometimes after a log-transformation. For the size of the individuals from the natural populations (Measure 4), we used a linear mixed-effect model (LMM) with sampling dates and ponds niched in the infection status and in egg status followed by Tukey contrast. Mobility was analyzed using a GLM with a Gamma error term and an inverse link function when the residuals were non-normal, each analysis being coupled with the two-sided Tukey contrast for pairwise comparisons. Concerning *D. magna*’s coloration (Measure 6), we found three peaks that were compared between non-white and white individuals using Wilcoxon signed-rank tests as data were not normally distributed. The global difference between the two spectra was not statistically tested (i.e., only a visual analysis).

We compared search and handling times (Measure 7) by *Notonecta* between infected (white) and uninfected *D. magna* (both for each of the three prey separately and with pooled prey) using paired two-sample one-sided t-tests when the data were normally distributed and paired one-sided Wilcoxon signed-rank tests when they were not. The values of Manly’s alpha index (Measure 8) were compared to the theoretical value of 0.5 indicating no prey choice using a one-sided t-test to detect a significant preference for infected over healthy *D. magna*.

We finally estimated a value of prey profitability for *D. magna* from the La Villette pond, in mJ/s, using the ratio between the total energy content (in mJ/*Daphnia*) and the handling time by *Notonecta* sp. for both healthy and infected *D. magna*. Based on the data obtained (Measures 5 and 7), 100 healthy and 100 infected *D. magna* were generated using a bootstrapped method (5,000 iterations). This procedure allowed computing a profitability for each individual. According to the bootstrap method, the 95% confidence interval of prey profitability is delimited by the 2.5% and 97.5% percentiles of the mean profitability distribution. We also, for each iteration, tested the effect of the infection on the predicted profitabilities using Wilcoxon signed-rank tests. We compared the distribution of these p-values to the distribution of p-values calculated from tests on randomized profitabilities (i.e., as a null model), and to a uniform distribution (Bland, 2013) with a Kolmogorov-Smirnov test.

## Results

### Experimental infection (Measures 1, 2, 3, and 4)

The three groups of *Daphnia magna*: control, infected and exposed, are phenotypically different (Fig. 1). We can observe that the ellipses of the 95% interval confidence of the means do not overlap (Fig. 1b). To summarize, Control individuals have either a long lifespan and intermediate mobility or high mobility and intermediate lifespan. Exposed individuals are close to the Control but with lower mobility and intermediate lifespan. Infected individuals show lower lifespan and fitness (total egg production), larger body size and varying mobility. Results are similar for the natural populations (Appendix B), with no effect on fecundity, lower mobility and higher body size for infected individuals. In detail, Axes 1 and 2 of the MFA (30% and 21.3% of the total variation) allow us to separate the three *D. magna* groups while the Control and Exposed groups and not distinguishable according to Axis 3 (16% of the total variation). Axis 1 represents Lifespan (positive correlation, p-value < 0.001) and Size (negative correlation, p-value < 0.001, Fig. 1a, 1c). Note that total egg production is mainly correlated to lifespan, rather than fecundity parameters. Axis 1 allows separating infected individuals that have a lower lifespan and a larger size, but a lower egg production, leading to a negative correlation between lifespan-egg production and si ze. Axis 2 corresponds to Mobility (negative correlation, p-values < 0.001 for four parameters). Fecundity can be described by Axes 1 and 2 as follows: Age at maturity and Clutch size are, respectively, positively (p-value < 0.001) and negatively (p-value = 0.009) correlated to Axis 1 while Clutch Frequency (p-value = 0.022) and Clutch Size (p-value < 0.001) are negatively correlated to Axis 2. Axis 1 is therefore sufficient to separate Infected individuals from the others, although both Axes 1 and 2 are necessary to separate Control and Exposed individuals.

**Figure 1.**
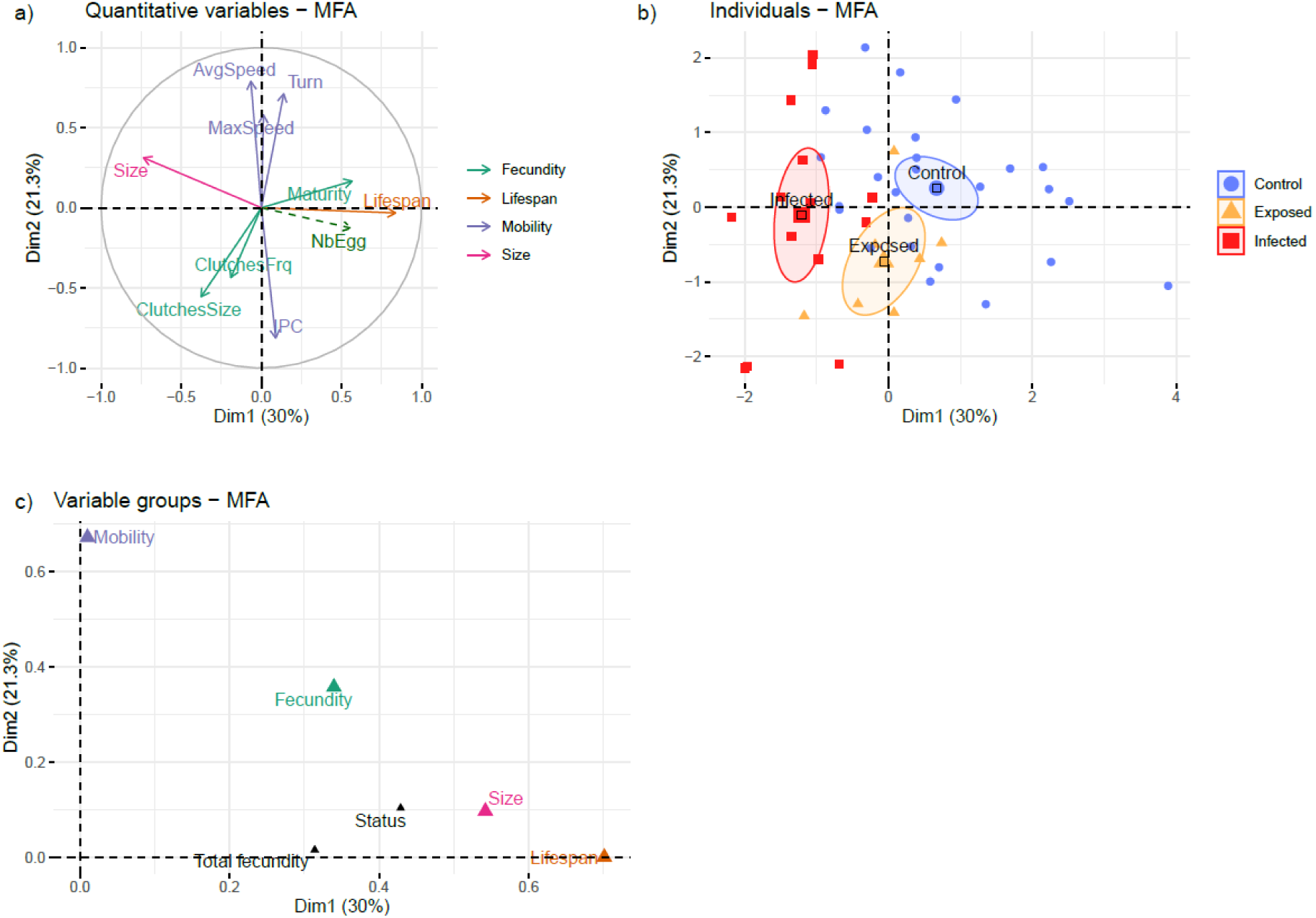
MFA on measurements of *D. magna* experimentally infected with DIV-1 for the two first dimensions. a) Quantitative variables grouped in four categories; note that total egg production (NbEgg) is a supplementary variable. b) Representation of individuals with ellipses for the 95% confidence interval. c) Representation of the group for the two dimensions.

### Biochemical composition and energy value (Measure 5)

We observed similar patterns in the two ponds sampled (p-values (status x pond) > 0.3, Table 2 and Fig. C3). Naturally-infected (i.e., white) individuals of *D. magna* had more proteins than healthy (i.e., non-white) specimens (p-value < 0.001 for La Villette), but the same amount of proteins *per* mg of *D. magna* as healthy brooding *D. magna* (p-value = 0.275 for La Villette). Infection and brooding did not change the amount of triglycerides whereas carbohydrates are increased in the presence of eggs/embryos alone (p-values < 0.001). To conclude, brooding and infected *D. magna* had a higher energy content if we consider both the energy *per* mg of *D. magna* and the energy *per* individual (all p-values < 0.003).

**Table 2.**
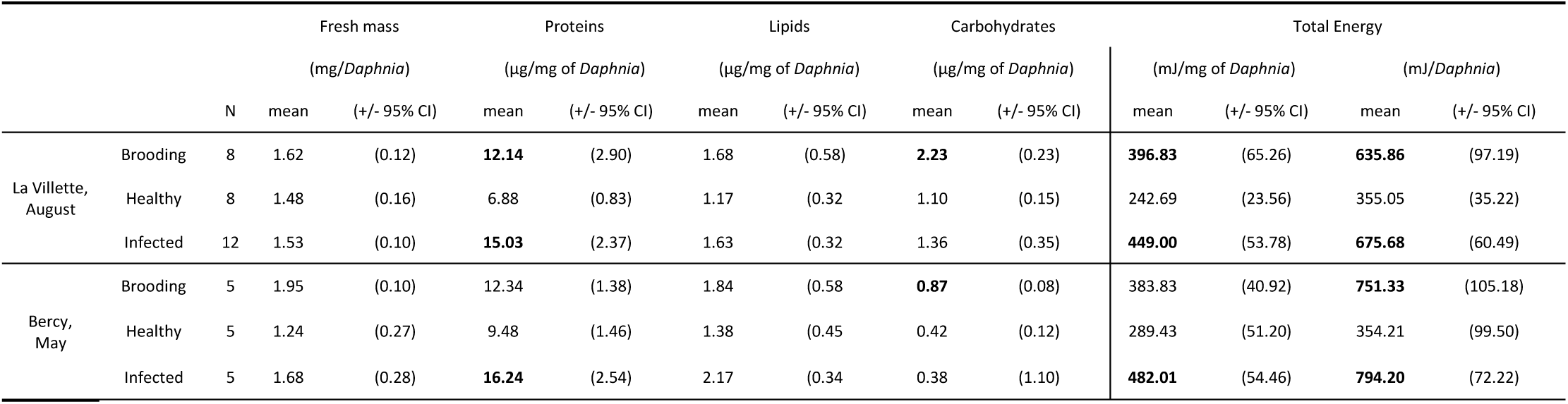
Host biomass and biochemical composition for the two populations. Means in bold are significantly different at 5% from healthy *D. magna*. See Table C5 for statistical values.

### Reflectance (Measure 6)

*Daphnia magna*’s reflectance (Fig. 2), measured as the percentage of reflected light (i.e., the more the light is reflected, the more the individual is colored for each wavelength/color, of white *D. magna* (likely infected) clearly shows that the white phenotype is associated with increased coloration (intensity) both in the UV and visible domains, and to a lesser extent in the infrared (280 to 850 nm). The overall reflectance of white *D. magna* was higher (12.19 +/-4.76%) than that of non-white *D. magna* (3.88 +/-1.47%). Three peaks of reflectanc e were observed for non-white *D. magna*: a first in UV around 317 nm, a second in blue around 460 nm, and a third in orange around 588 nm. Few differences were observed on the position of the three peaks of reflectance in white *D. magna* with a small shift toward green for the blue and orange peaks (around 477 and 570 nm, respectively; p-values < 0.001), but at the same position for the UV peaks (around 314 nm, p-value = 0.083).

**Figure 2.**
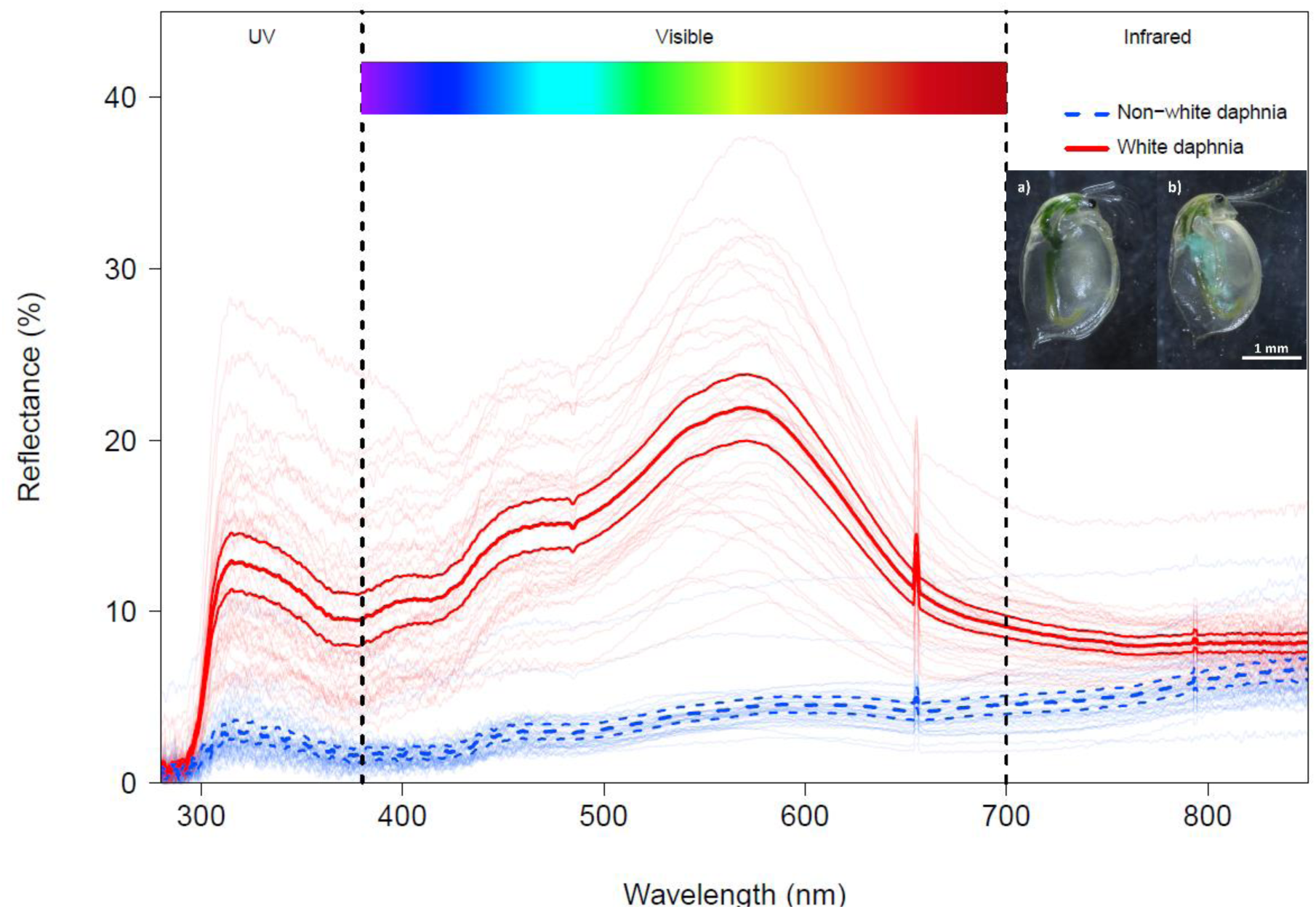
Effects of DIV-1 on reflectance between 280 and 850 nm. Blue (dashed) lines are healthy *D. magna* and red (solid) lines are infected *D. magna*. Highly visible lines are the mean and the lower and upper 95% confidence interval. Weakly visible lines correspond to all the measured *D. magna*. Note the two peaks due to the material (artefacts) around 660 nm and 790 nm. See Table C6 for statistical values. On photographies, a) is non-white daphnia and b) is white daphnia (from Prosnier, 2018), see also Fig C1.

### Vulnerability to predation (Measures 7, 8, and 9)

For both predator species, the time elapsed between two consecutive captures (Measure 7) did not differ between naturally-infected (i.e., white) and uninfected (i.e., non-white) *D. magna* (Fig. 3a, Fig. A1). However, the handling time by *Notonecta* was significantly longer when they consumed infected *D. magna* (p-value <0.001 for all catches, Fig. 3b), which are also preferred (Measure 8) over healthy *D. magna* (p-value = 0.03, Fig. 3c).

**Figure 3.**
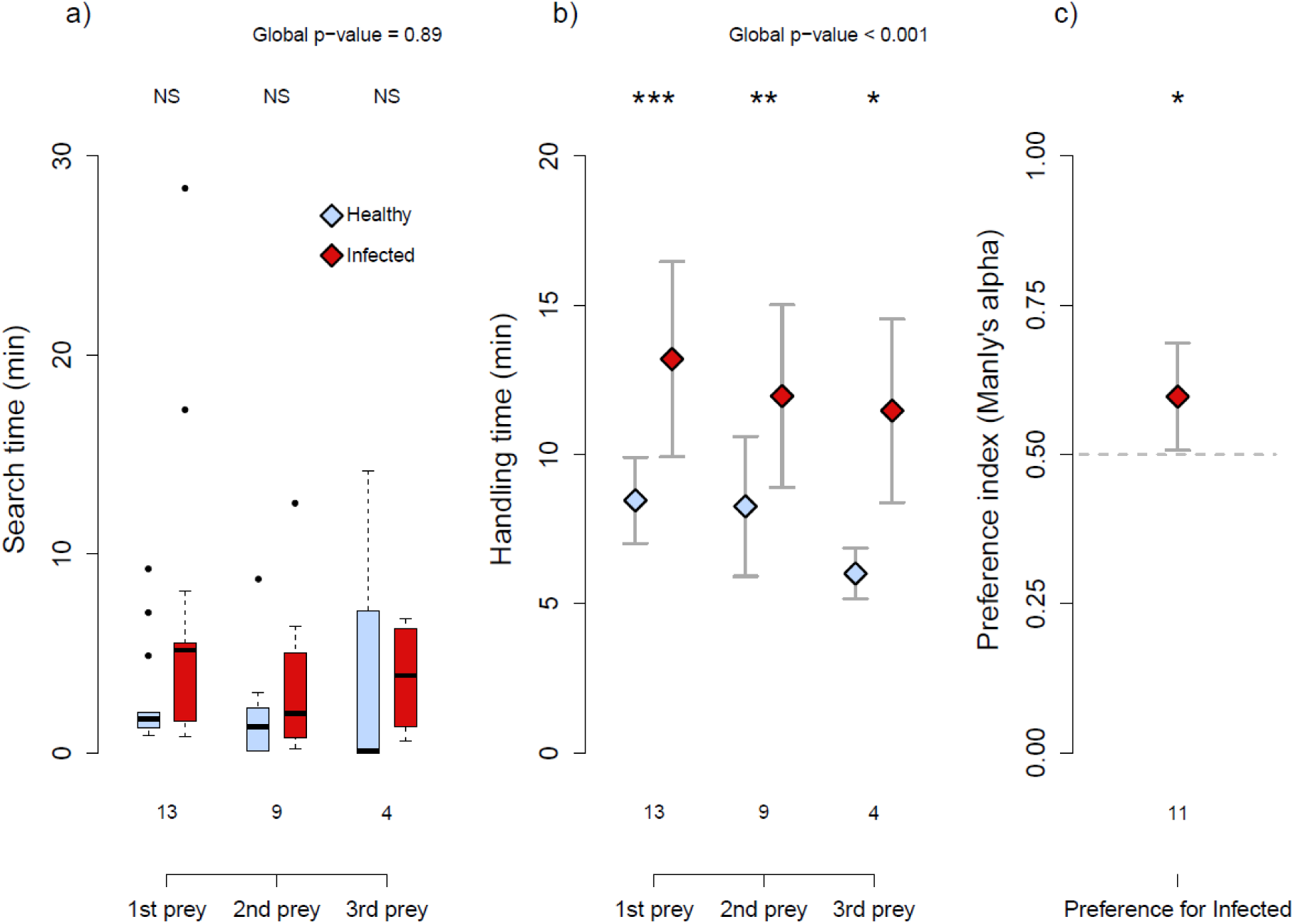
Effects of DIV-1 on vulnerability to predation. a) Search time and b) handling time by *Notonecta* sp., healthy (light blue) or infected (dark red), for the three prey; c) preference for infected *D. magna*. a,b) Statistics compare healthy versus infected prey: dot P < 0.1, *P < 0.05; **P < 0.01; ***P < 0.001; NS P>0.1. a) Central bars represent the median, boxes the interquartile range, and dots the outliers (> 1.5 times the interquartile range); b,c) diamonds represent the means and bars the 95% confidence intervals. See Table C7 for statistical values.

### Prey profitability

Using the values of handling time (Measure 7) and total energy content *per D. magna* (Measure 5), we determined profitability with a bootstrap analysis. The profitability of healthy *D. magna* is 49.51 mJ/ind. (95% CI: 49.98 – 57.66) and that of infected *D. magna* is 62.19 mJ/ind. (95% CI: 57.97 – 68.09). Following Cumming & Finch (2005) about the non-superposition of the 95% confidence intervals and based on the p-value distributions, the profitability of naturally-infected (i.e., white) *D. magna* is significantly higher than the profitability of healthy (i.e., non-white) ones. This is confirmed by the analysis of the distribution of p-values for the effect of infection, that is significantly different from the null model and from the uniform distribution (p-values < 0.001, Table C8). Note that the null model is not different from the uniform distribution (p-value = 0.347) as expected (Bland, 2013).

## Discussion

Parasites may affect their host in many ways, with potential repercussions on their predators. Here, we investigated the direct and indirect effects of iridovirus DIV-1 (Daphnia iridescent virus 1) infection on *D. magna* water fleas. Note that we considered along this study that individuals with the white phenotype (i.e., previously named the White Fat Cell Disease) are infected by DIV-1 (Toenshoff et al., 2018), and that non-white individuals are not infected (but see the discussion about exposed individuals from the experimental infection). We found that DIV-1 reduced the survival of water fleas, while the effects on fecundity were not significant. We also observed that infection changed the phenotype of *Daphnia*, mainly by increasing host size, coloration, and energy content. About *Notonecta* predation, infection increase handling time but not affect search time. As a result, the profitability of infected individuals was increased by 24%. Based on the optimal foraging theory, a preference for infected individuals should be expected, and this assumption is supported by our results. We will after discuss the specific characteristics of “exposed individuals”, those experimentally presented to the virus but displaying no visible sign of infection (white coloration). Finally, we will highlight the complex consequences of parasitism on trophic links.

### Reduction of survival but limited effects on vulnerability to predation

The stronger effect of infection concerns the reduction in *D. magna* lifespan. However, there is no obvious effect on fecundity: no change in clutch size or clutch frequency, contrary to previous affirmation of a lower fecundity in the same host-parasite system (Ebert, 2005). The only modification in terms of fecundity characteristics was the earlier age at maturity, as previously reported with *D. magna* infected by a microsporidian (Chadwick & Little, 2005). This change could be a plastic modification to compensate for the shorter lifespan (Agnew et al., 1999). Despite this compensation, the total number of offspring was lower for infected *D. magna* compared to control *D. magna*, thus illustrating the negative effect of infection on fitness. In support of our finding, this virulence effect was already observed by Ebert et al. (2000) and Decaestecker et al. (2003) who reported an effect on lifespan and total number of offspring, although these authors did not analyze the effects on clutch size or fecundity. Due to the virus replication and accumulation (Marina et al., 2003; Toenshoff et al., 2018), host physiology and integrity are expected to be largely impaired (Agnew et al., 1999). DIV-1 thus reduced host fitness (i.e., total offspring produced during lifetime) by increasing direct adult mortality, which likely contributes to explain its low prevalence in ponds (Decaestecker et al., 2005). No effect on juvenile mortality was observed due to the virus exposure, which supports the previous hypothesis (Agnew et al., 1999; Marina et al., 2003; Toenshoff et al., 2018) that the virus progressively accumulates inside the host and ultimately leads to death.

Altered body size, mobility, coloration, and biochemical content, could lead to indirect effects through the modification of trophic interactions. DIV-1 infected individuals are larger, a change that is generally observed in case of infection by castrating parasites (Hall et al., 2007), where the energy not used to reproduce is reallocated to growth. Here, there is no effect on fecundity, meaning that an unknown physiological modification could explain it. A possible explanation would be that lower speeds (higher speeds being generally associated with larger sizes, see Dodson & Ramcharan, 1991) save part of the individual’s energy budget that could be reinvested in somatic growth. The difference between ponds in terms of speed and carbohydrate content may be due to differences in the genotypes of both DIV-1 and *D. magna*, as virulence is known to vary with genotypes (Decaestecker et al., 2003). This hypothesis should be tested with experimental infestations for the two populations and also with cross-infestations – combined with genotype analysis. Abiotic conditions may also determine how hosts deal with infection (Bedhomme et al., 2004) and biotic pressure due to predation. We only found predators of *Daphnia* sp. (Chaoboridae) in the La Villette pond (pers. obs.) where *D. magna* are less active. Because Chaoboridae larvae are ambush predators (Spitze, 1985), fast *D. magna* might encounter more predators and thus be more prone to predation (Gerritsen & Strickler, 1977), leading to the lower speed of this *D. magna* population. The presence of a predator could also affect other phenotypic characteristic as body size: larger individuals in presence of *Chaoborus* but smaller individuals in presence of fish (Riessen, 1999). As a result, this would mask the differences between healthy and infected individuals. Other works have shown that the speed of *Daphnia* sp. could affect vulnerability to predation with slow Cladocera being more vulnerable to copepods (Chang & Hanazato, 2003) and fish (O’Keefe et al., 1998). Moreover, due to the structural properties of iridovirus causing iridescence (Williams, 2008), infected *D. magna* showed a higher reflectance in the UV and visible domains than *D. magna* presenting no sign of infection (i.e., non-white). Infected *D. magna* may thus become more visible (especially considering the larger size of infected individuals) and more attractive (O’Keefe et al., 1998; Modarressie et al., 2013; Jacquin et al., 2013) for *Notonecta* sp., which has a high sensitivity in UV (375 nm) and green (520 nm) (Bennett & Ruck, 1970). This is consistent with the observed preference of *Notonecta* sp. for infected *D. magna*. It would be interesting to determine the relative importance of the various phenotypic changes observed in infected individuals, whether predators prefer infected individuals because they are larger, slower, more visible, or due to the changes in energy contents.

### Increase in host energy content leads to higher profitability

Because of the parasite’s requirements and the host’s immune response, infection is likely to alter the biochemical composition of the host. For instance, the fungi *Polycaryum laeve* reduces the lipid content of their *Daphnia pulicaria* hosts (Forshay et al., 2008), while infection by *Polymorphus minutus* (acanthocephalan) increases the triglyceride content of *Gammarus roeseli* (Médoc et al., 2011). Here, we showed that the energy content of DIV-1-infected *D. magna* is higher than that of broodless healthy ones but comparable to that of healthy individuals with eggs, illustrating how the effects can change with parasite species. The difference in biochemical composition between infected and uninfected *D. magna* is due to variations in the protein contents and makes that infected *D. magna* could be more nutritious. This could be linked to the virus’ life cycle that uses the cellular machinery of the host to produce the viral proteins of their capsids with the persistence of the virus in *D. magna* until host death. An alternative explanation of the higher protein content could be the immune response of the host that would use antimicrobial peptides (McTaggart et al., 2009; Rosa & Barracco, 2010; Xie et al., 2016). Although the fat cells of DIV-1-infected *D. magna* are described as being larger by Toenshoff et al. (2018), we found no difference in the lipid content between infected and uninfected *D. magna*. Overall, except for the carbohydrates, the biochemical composition of infected *D. magna* was closer to that of brooding *D. magna* compared to uninfected *D. magna*. This effect is magnified by the larger size of infected individuals, leading to the higher energy content of infected *D. magna*.

Optimal foraging theory predicts that predators should maximize net energy gain (MacArthur & Pianka, 1966; Charnov, 1976a; b). Following our estimations of *D. magna* energy content and handling time by *Notonecta* sp., we approximated *D. magna* profitability to be around 50 and 62 mJ/s for uninfected and infected individuals, respectively, representing an increase of 24%. Consequently, in spite of a higher handling time, possibly due to the fact that the prey are bigger, the large increase in energy content leads to a higher profitability for infected individuals. Search time, the third parameter of net energy gain is unchanged despite the modifications to host coloration and a possible reduction in mobility (also in the preliminary experiment with fish). Consequently, based on search time, handling time, and energy content, the predator’s preference for infected *D. magna* is not surprising. Nevertheless, we also showed that the parasite greatly increased host mortality, probably leading to the low prevalence observed in natural populations (0.5-3%). Thus, high virulence could counterbalance the increase in host profitability, limiting predation rate on infected prey under natural conditions. In addition, the low prevalence may explain why the meta-analysis of Flick et al. (2016) showed that predators rarely modify their preference for infected prey. Long-term experiments with predators of *Daphnia* while controlling DIV-1 prevalence to dampen the direct effects of DIV-1 infection could be undertaken to explore the indirect effects of parasites on predators’ diet.

### Exposed individuals differ from healthy ones

Some individuals were exposed to DIV-1 but did not exhibit the most visible sign of virus infection, namely, the white coloration. Nevertheless, we noted two differences between these so called “exposed” individuals and healthy individuals: a lower lifespan and a lower mobility. We propose three hypotheses to explain these differences. First, they could have escape infection. Results on healthy *D. magna* showed that their lower mobility is positively correlated with a longer lifespan. Therefore, if exposed individuals have escaped infection, for instance because they are slower and thus encounter the virus less often, then they should have a longer lifespan. However, because exposed *D. magna* have a shorter lifespan, we may suppose that they have been infected by the virus and not only escaped infection. More, due to our setup where microcosms are small and the medium daily resuspended, this escaped explanation seems unlikely. Second, they could have resisted to infection. Compared to *D. magna* displaying the white phenotype, we observe that this resistance results in a lower lifespan reduction (probably because the virus does not accumulate in the host) but in a greater mobility reduction. Both effects may occur because resistance (immunity) is energetically costly. Dallas et al. (2016) showed the “cost of resistance” (lifespan reduction) on various *Daphnia* sp. exposed to *Metschnikowia bicuspidata* (fungi). On the contrary, Labbé et al. (2010), with their experiment of *D. magna* infected by the bacteria *Pasteuria ramosa*, did not observe such costs. However, the between-species comparison remains limited as the cost of resistance should depend on the immunity system, which differs between fungi, bacterial and virus infection (McTaggart et al., 2009). A third hypothesis is that DIV-1 effectively infects these specimens of *D. magna* without inducing the white phenotype. Studies on iridovirus called this effect “covert infection” in opposition to “patent infection” (Williams, 1993; Marina et al., 1999; Williams et al., 2005). We conclude from these observations that there are not two extreme categories (i.e., healthy versus infected) with an intensity gradient of parasitic effects but rather various combinations of effects depending on how the host react to infection. Clarifying this aspect would require testing whether exposed individuals are infected or not using microscopy or PCR techniques (Toenshoff et al., 2018). Thus, in the continuity of this study, we question how this third category is important in *D. magna* populations, how they are affected in terms of energy content, and thus what are their consequences in terms of predator diet and at larger scales.

### On the complexity of adding parasites to predator-prey relationships

In addition to the well-known virulence effect (i.e., higher mortality) leading to reduced abundance, we showed some less studied morphological, behavioral, and physiological alterations resulting in increased profitability. Thus, at larger ecological scales, two opposite effects could be expected considering the optimal foraging theory. The increase in profitability should promote the preference of predators whereas reduced availability due to the higher mortality should decrease encounters and thus the inclusion of infected *D. magna* in the diet. While the evolutionary investigations of the consequences of prey infection on predator’s diet go beyond the scope of the present article, theory suggests antagonistic effects between increased host vulnerability, which should favor predation on the host, and increased host mortality, which acts in the opposite way (Prosnier et al., 2020). It would be interesting to perform experiments with and without infection dynamics, that is, by fixing or not host density or parasite prevalence to separately consider the effects on host energy and host availability. Such experiments would also offer a way to understand how predation on host affects parasite dynamic, the conditions under which it reduces infection (healthy herd hypothesis, Packer et al., 2003) or when it favors the dispersal of a parasite that is not transmitted trophically, as *Chaoborus* do for the spores of a fungal parasite of *D. dentifera* (Cáceres et al., 2009).

A second interesting point is the existence of a more complex structure in the host population with the exposed individuals showing cryptic phenotypes (covert infection). They are rarely studied in experimental work (partly due to the difficulty in identifying them) despite their likely high prevalence compared to individuals with visible signs of infection (Marina et al., 1999; Williams et al., 2005). For instance, here, we found a very low prevalence of DIV-1 (3%) based on individuals showing the white phenotype, suggesting little consequence on ecological dynamics. However, if there is a high prevalence of covert-infected *D. magna* showing (at least) reduced survival and mobility, then consequences on communities should be stronger than expected from the prevalence and phenotype alterations of patent-infected individuals only. Covert infection could explain why our apparently “healthy” individuals are more variable in terms of mobility than the infected ones, with potentially bigger differences between *D. magna* that are actually uninfected and patent-infected individuals. In theoretical work, there are interesting studies on various epidemiological models (like SEIR) that could be adapted by taking into account the additional category of exposed individuals (e.g., Sorrell et al., 2009; Britton & Jane White, 2021).

To conclude, we encourage further studies at a larger ecological scale, considering that prey infection has repercussions on predators (Flick et al., 2016) and other potential prey (Decaestecker et al., 2015; Prosnier et al., 2018), potentially leading to the modification of trophic links. As shown in many food web studies, it is crucial to understand the implications of parasites on community composition, stability, and functioning (McCann, 2000; Kondoh, 2003; Frainer et al., 2018).

## Acknowledgment

The authors would like to thank all the people who contributed to the success of these experiments: Tiphaine Boursier, Baptiste Carrere, and Léo Bricout for performing the preliminary experiments; Pierre Fédérici, Jérôme Mathieu, Gérard Lacroix, Thomas Tully, Thibaud Monnin, Gabrielle Ringot, Julien Gasparini, Adrien Frantz, Clotilde Biard, and Eric Edeline for lending material and providing help; and Claude Yéprémian for providing the *Scenedesmus* sp. and Lauren Boutier for providing the European minnows. The authors also thank Victoria Grace for reviewing the English language, and David Civitello for a useful comment. Finally, the authors thank Luis Schiesari, Thierry De Meeus, and Eglantine Mathieu-Bégné for their useful comments.

## Data, script and code availability

Data, script and code are available on Zenodo. DOI: 10.5281/zenodo.6006617 (Prosnier et al., 2022)

## Supplementary information

Supplementary information is available after the references:

Appendix A: Vulnerability to fish predation

Appendix B: Comparative analysis for experimentally and naturally-infected individuals

Appendix C: Supplementary figure and tables of statistics

## Appendix A: Vulnerability to fish predation

We did not observe any effect of DIV-1 infection on the intercapture time of *Daphnia magna* by *Notonecta* sp. despite the color modification. Thus, in line with our hypothesis, we tested whether it could affect the intercapture time of an aquatic vertebrate: the European minnow (*Phoxinus phoxinus*). Using another predator that varies in terms of size, mobility, vision, and hunting method is more representative of the diversity of strategies used by predators of *D. magna* in the field.

Fish (2.6-3.4 cm in total length) were purchased online (Armorvif, Brittany, France) and kept in a rearing room under natural light at 19 °C, at a density of 1.7 fish.L^-1^. The water comprised 75% of spring water (Cristaline®, Cristal-Roc source) and 25% of osmotic water, which was regularly changed (>30% volume per week) and cleaned daily with a net. The fish were fed with commercial food pellets (Goldfish premium, Tetra®), twice a week.

Fish (n = 46) were starved for at least 24 h before the experiments to standardize predation. The experiments were performed in an aquarium (34×19×24cm) filled with 10 L of water (75% spring water, Cristaline®, Cristal-Roc source, and 25% osmotic water). To be close to the visual environment of the animals, we covered the edges of the aquarium with green plastic and the bottom with brown paper. The length of the aquarium was divided into two equal parts with a central wall made of green plastic: one part of the aquarium contained the fish and the other part three infected or uninfected *D. magna* without eggs. After a 1-h acclimation period, we removed the central wall to start the experiment with the fish being allowed to forage for 1 h. Predation events were recorded with a webcam (Logitech HD Webcam Pro C920) and the software OBS Studio (version 21.1.2). We measured the time of each capture and thus obtained the duration between the predation events (first, second, and third capture). Each fish experienced the two different types of prey with 1 h between the two experiments. To avoid time and order effect, half of the fish started with healthy *D. magna* and the others with infected *D. magna*. After 1 h, we performed the same experiments with the other prey type *per* predator.

We compared fish’s search time between the two prey types using paired two-sample t-tests after a log-transformation, leading to normally distributed and homoscedastic data. Despite the lower search time for the first prey (Fig. A1, Table C7, p-value = 0.04), we did not observe any effect for the second and third prey (p-values > 0.44). This suggests that the possible effects of DIV-1 infection on coloration and mobility did not influence the search time of the European minnow. Contrary to the predation tests made with *Notonecta*, body size was the same between infected and uninfected *D. magna* (p-value = 0.803).

**Figure A1.**
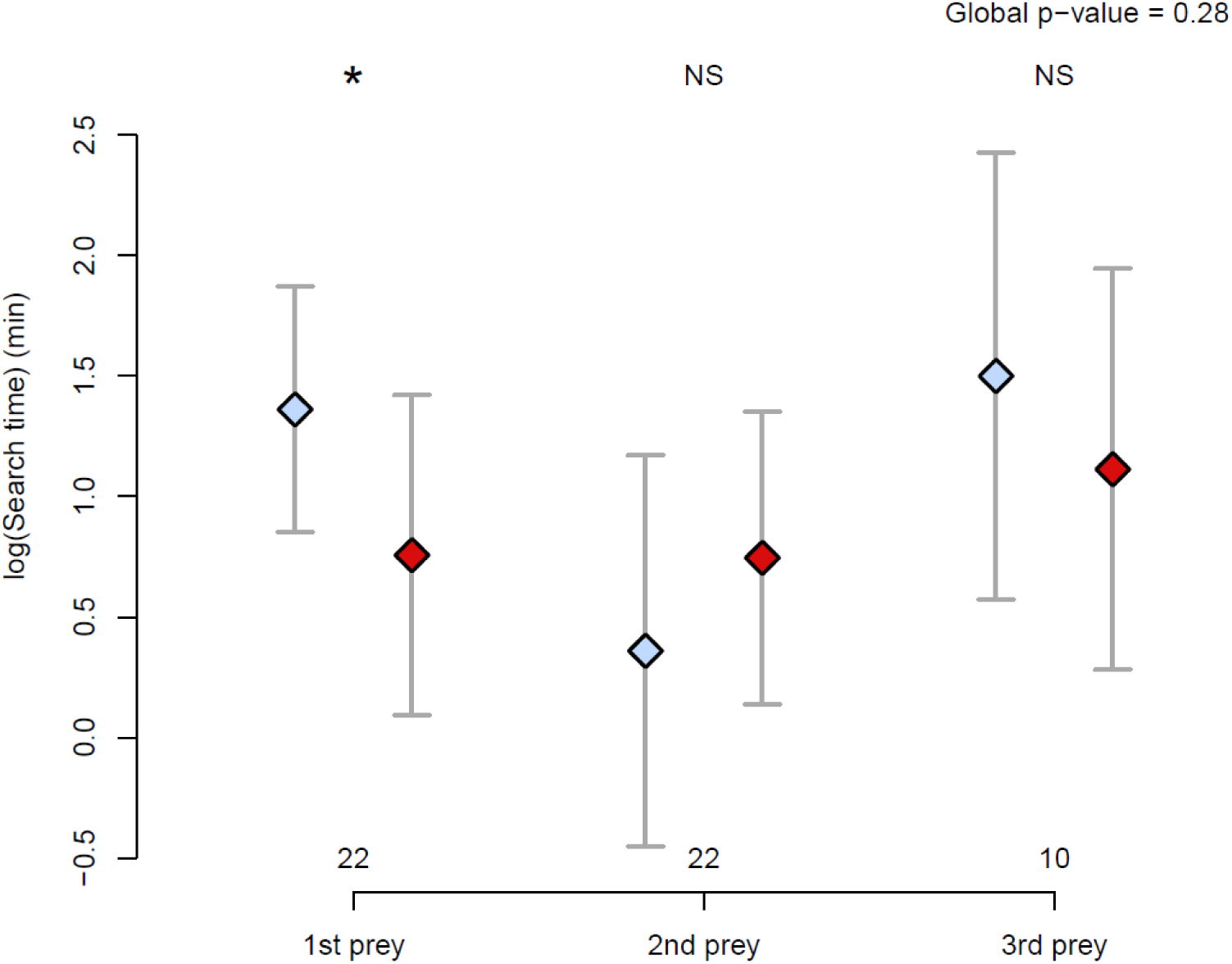
Effects of DIV-1 infection on vulnerability to predation by fish. Search time on healthy (light blue) or infected (dark red) prey for the three prey. Statistics compare healthy versus infected prey: dot P < 0.1, *P < 0.05; **P < 0.01; NS P>0.1. Dots represent the means and bars the 95% confidence intervals. See Table A2 for statistical values. See Table C7 for statistical values.

## Appendix B: Compared analysis of *Daphnia magna* traits for both experimental and natural infection

### Methods

#### Fecundity (Measure 1)

For naturally-infected individuals, collection took place in April-June 2018 in the two ponds (Bercy and La Villette). We sampled 20 L of water filtered with a 50 µm net to collect D. magna. After sorting white and non-white D. magna, individuals were fixed using glycerol solution (1% glycerol, 70% ethanol, 29% water). We then categorized individuals as broodless (without eggs nor ephippia), egg-carrying (with parthenogenetic eggs), and ephippia-carrying (with sexual ephippia).

#### Mobility (Measure 3)

Naturally-infected individuals were collected from the La Villette pond in September 2017 (n = 188) and the Bercy pond in May 2018 (n = 135), stored in rearing tanks and processed within a day after collection. Mobility was measured as for experimentally-infected individuals (see the Matherial and Methods in the main text).

#### Body size (Measure 4)

Body size of naturally-infected individuals, among those collected in the La Villette and Bercy ponds (Measure 1, n = 435), was measured with a micrometer screw.

#### Statistical analysis

In addition to the MFA, we performed a survival analysis on the results of experimental infections (log-rank test) and compared the death age between healthy juveniles (control *D. magna* dead before the first clutch) and exposed juveniles to assess juvenile mortality (Measure 2). For adult mortality (from first clutch to death), we compared the death age (i.e., the survival) between healthy (control), exposed (no white phenotype), and infected *D. magna* (with the white phenotype) and the adult period (from first clutch to death). To quantify the effects on reproduction (Measure 1), we performed a survival analysis (log-rank test) on age at maturity (date of the first clutch) and compared clutch frequency and mean clutch size (i.e., number of eggs/embryos in the brood chamber) between adult categories using a generalized linear mixed-effect model (GLMM) with a Gamma error term and an inverse link function, and mother (i.e., clonal lineage) as random effect, followed by one-sided Tukey contrast for pairwise analyses. Total reproduction (total number of clutches and offspring during lifetime) was analyzed using a GLMM with a Poisson error term and a logarithmic link function, and mother (i.e., clonal lineage) as random effect, while we used one-sided Tukey contrast for pairwise analyses. See a summary in Fig. C2.

To analyze the fecundity of naturally-infected individuals (Measure 1), we considered the abundances of broodless (no egg nor ephippia), egg-carrying, and ephippia-carrying *D. magna* with (i.e., infected) or without (i.e., healthy) phenotypic signs of infection. Because infection is visible around Day 10, we considered all infected *D. magna* as adults. However, a large proportion of broodless healthy *D. magna* could be juveniles (Hülsmann & Weiler, 2000). Thus, using the Lampert’s method (described in Stibor & Lampert, 1993) considering as adult size the smallest class size where less than 50% are broodless, we determined adult size and thus the proportion of adults in each pond. We calculated the number of adults in the broodless group based on this proportion. With this correction, we expected to limit the overestimation of infected brooding *D. magna*. We compared the abundances of the infected and healthy groups with a Fisher’s exact test, because several groups showed a low abundance.

Analyses of mobility (Measure 3: average speed, maximal speed, proportion of inactivity time, number of turnings) and body size (Measure 4) of experimentally-infected individuals were performed with a linear mixed-effect model, with mother (i.e., clonal lineage) as random effect, followed by two-sided Tukey contrast for pairwise analyses. For the mobility of naturally-infected individuals, we performed an ANOVA followed by pairwise-t-test when normality and homoscedasticity was verified, and otherwise a GLM with a Gamma error term and an inverse link function. For the size of naturally-infected individuals (Measure 4), we used a linear mixed-effect model with sample dates and ponds niched in the infection status and in egg status, followed by Tukey contrast.

#### Fecundity and mortality (Measures 1 and 2)

Experimental infection (Measure 1; Fig B1 and Table C1) significantly reduced the survival (p-value < 0.001, Fig. B1a) and adult lifespan (p-value < 0.001) of *D. magna*. DIV-1-exposed individuals (i.e., exposed to the parasite but presenting no apparent sign of infection) exhibited intermediate lifespan and duration of adult life compared to the two other experimental groups. Exposure to parasites did not affect the mortality of immature *D. magna* (p-value = 0.319, Fig. B1a). Age at maturity (first clutch) was significantly lower in infected *D. magna* than in controls (p-value = 0.037, Fig. B1b). Exposed individuals were not different from infected and control individuals in terms of age at maturity (p-values > 0.25). No difference was found for the mean clutch size (p-value = 0.895, Fig. B1c) and clutch frequency (p-value = 0.508, Fig. B1d) between each of the groups. DIV-1 significantly reduced the total number of clutches (p-value = <0.001) with an intermediate value for exposed *D. magna*. Infection reduced total offspring production (p-value < 0.001, Fig. B1e-f) with an intermediate value for exposed *D. magna*.

For natural populations (Measure 1; Fig B2 and Table C2), after applying the correction to exclude juveniles using the Lampert’s method, we did not observe any effect on fecundity (egg and ephippia production) except for the specimens collected from the Bercy pond on 19 April, which were characterized by higher amounts of ephippia and a lower egg production for infected *D. magna* (p-value = 0.022), and for those collected from the La Villette pond on 17 May, which had a lower fecundity for infected *D. magna* (p-value = 0.008).

**Figure B1.**
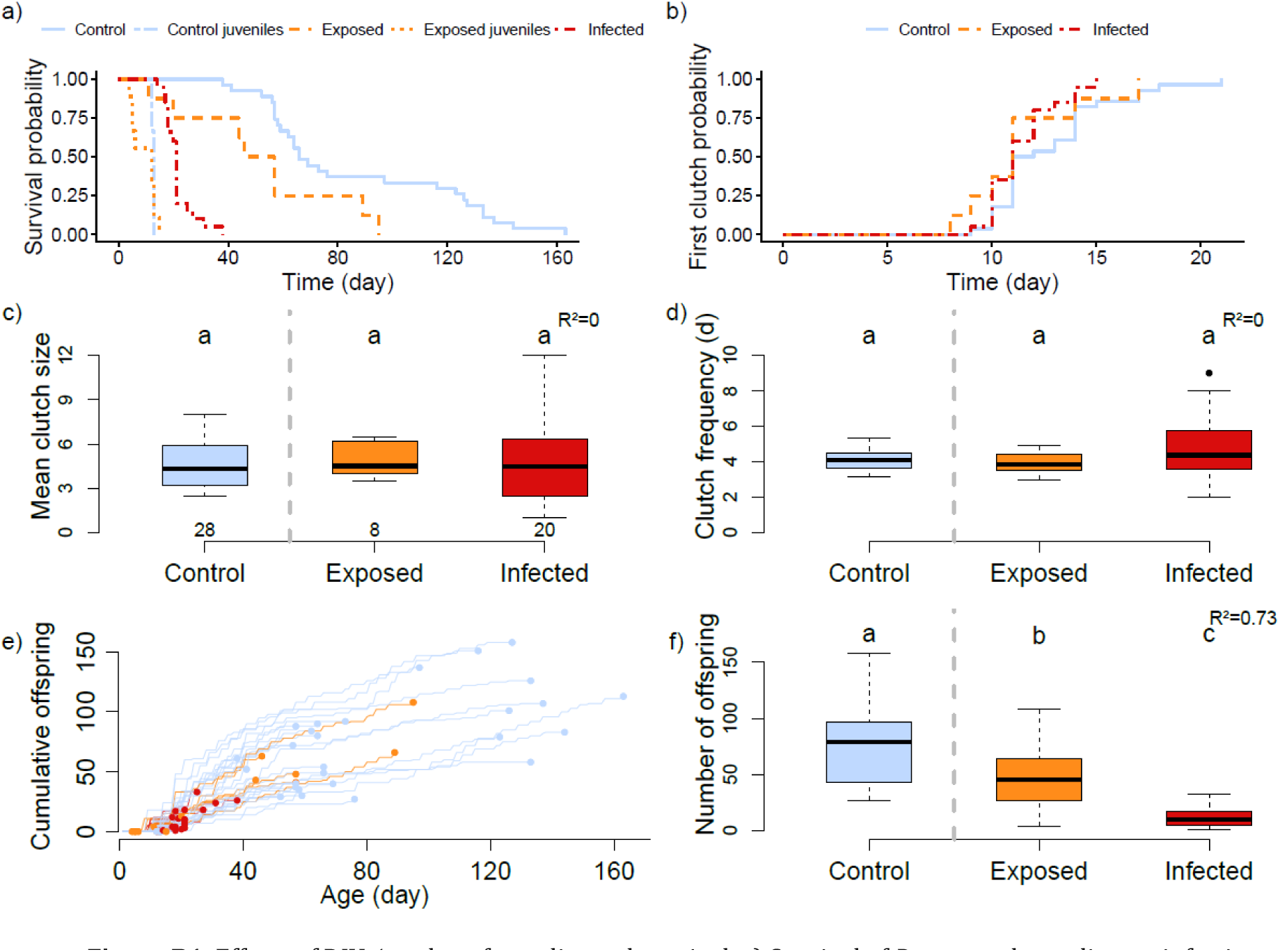
Effects of DIV-1 on host fecundity and survival. a) Survival of *D. magna* depending on infection status (healthy, exposed, or infected) and depending on whether or not they have offspring in their lifetime; b) age at maturity (first clutch); c) mean clutch size; d) clutch frequency; e) Cumulative offspring production per individual; and f) total number of offspring during lifetime for control, exposed, and infected *D. magna*. The vertical dashed line separates *D. magna* exposed to the control solution (left) and those exposed to the DIV-1 solution (right). Numbers in c) are the numbers of *D. magna* for each category. The same letters indicate the groups that are not significantly different at 0.05. a,b) Representation according to the Kaplan-Meier method; c,d,f) central bars represent the median, boxes the interquartile range, and dots the outliers (> 1.5 times the interquartile range); e) dots represent death of each individual, note that dots are the total number of offspring produced along life, thus are data of f). See Table C1 for statistical values.

#### Mobility (Measure 3)

For experimentally-infected *D. magna* (Fig. B3a, B3c, B3e, Table C3), exposed individuals showed lower activity with lower mean (p-value = 0.034) and maximum (p-value = 0.03) speeds, and were more often inactive (p-value = 0.029) than control individuals. Conversely, infected *D. magna* showed intermediate activity patterns, with no significant difference between healthy or infected individuals (p-values > 0.2). The number of turnings was higher for control *D. magna* compared to infected (p-value = 0.064) and exposed (p-value = 0.003) individuals. For naturally-infected *D. magna* (Fig. B3b, B3d, B3f), there was no significant difference in mobility between uninfected and infected *D. magna* from the La Villette pond, whereas infected *D. magna* from the Bercy pond compared to uninfected *D. magna* showed a significant decrease in mean and maximum speed, activity, and number of turnings (all p-values < 0.001). Note that we grouped healthy brooding and non-brooding *D. magna* together in the uninfected category, because eggs/embryos did not modify mobility (all p-values > 0.7).

**Figure B2.**
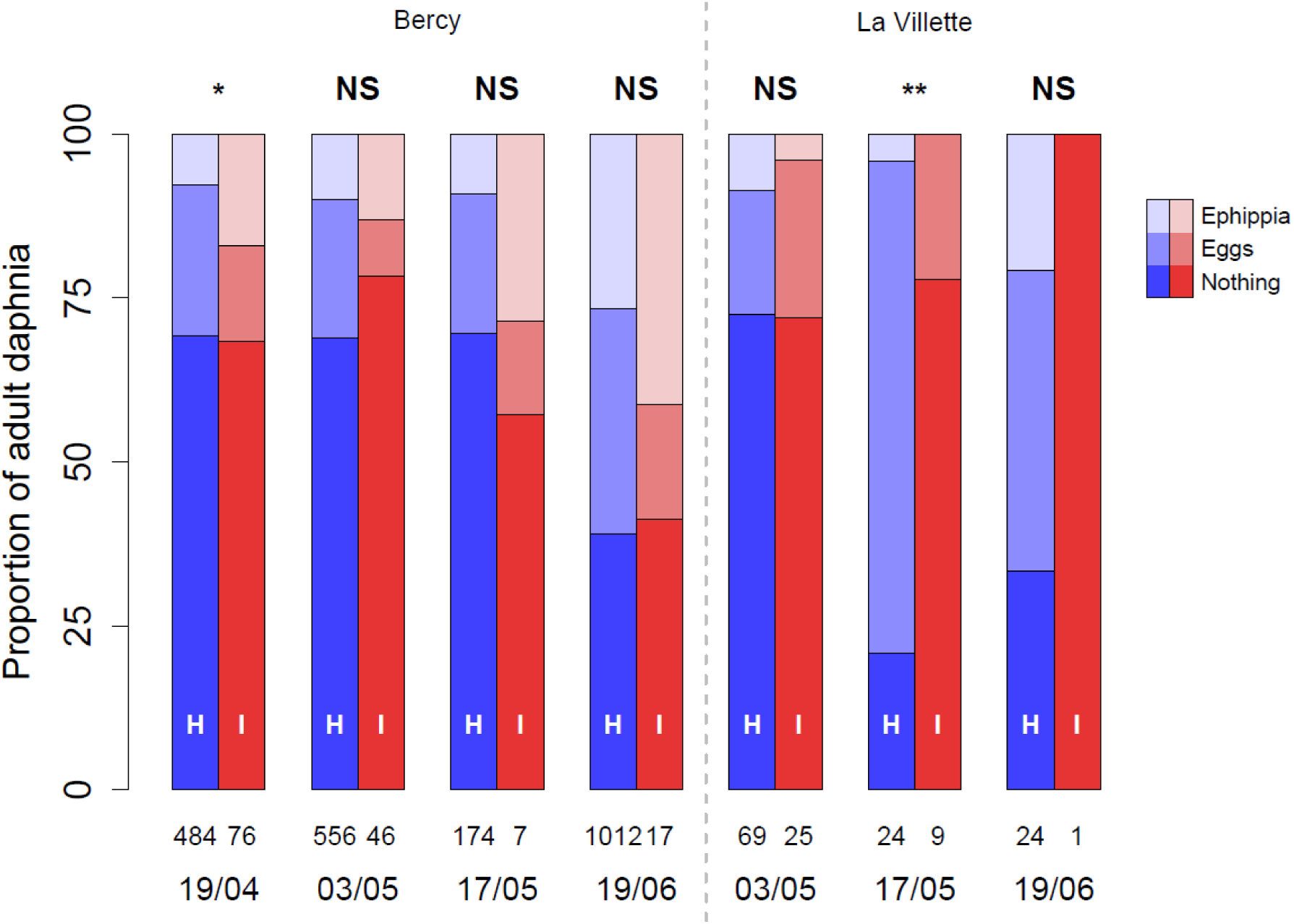
Proportion of adult *D. magna* without eggs, with eggs, or with ephippia depending on their infection status (healthy in blue, infected in red) in the two ponds for various dates. Numbers are the numbers of infected or uninfected *D. magna*. Statistics compare healthy versus infected prey: dot P < 0.1, *P < 0.05; **P < 0.01; NS P>0.1. See Table C2 for statistical values.

#### Body size (Measure 4)

We compared the size of healthy and infected *D. magna* (Fig. B4, Table C4). For experimentally-infected *D. magna* (same age), infected individuals were larger than controls (Fig. B4a, p-value = 0.033), while exposed *D. magna* had an intermediate size. For natural populations (Fig. B4b), we observed the largest sizes with infected individuals that were broodless (p-values = 0.01) but not with infected *D. magna* with eggs or ephippia (p-value > 0.22). Finally, for the two groups of naturally-infected individuals used for the predation experiments, only the infected *D. magna* used as prey for the *Notonecta* sp. were larger than healthy individuals (p-value < 0.001).

**Figure B3.**
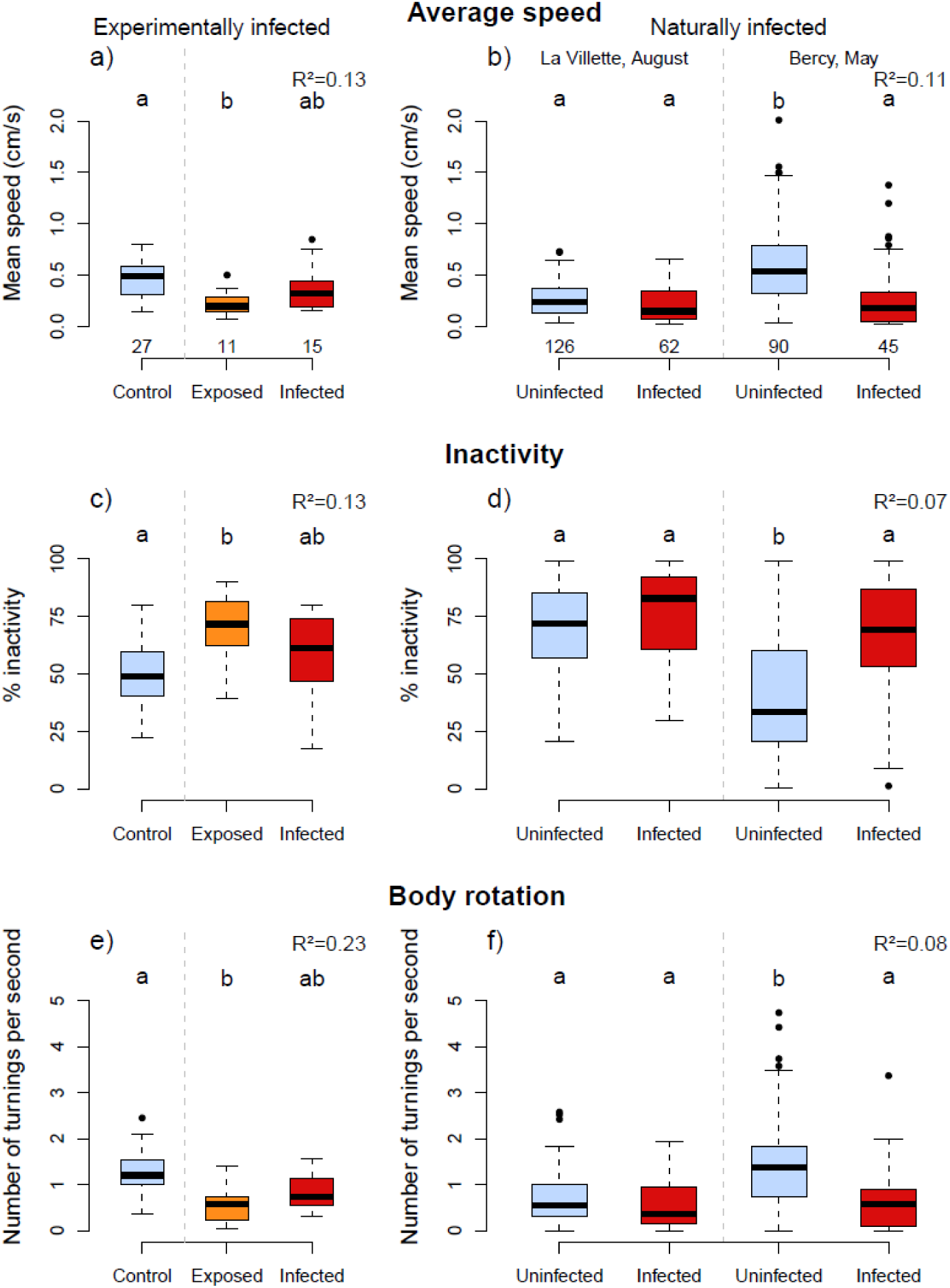
Effects of DIV-1 on host mobility on experimentally infected (left) and naturally infected (right) *D. magna*. a-b) Mean speed; c-d) proportion of inactive time; and e-f) number of turnings for *D. magna* with or without signs of DIV-1 infection. Note that the uninfected category aggregates brooding and unbrooding *D. magna,* because there was no statistical difference in their mobility. Numbers in a-b) are the numbers of *D. magna* for each category. The same letters indicate groups that are not significantly different at 0.05. Central bars represent the median, boxes the interquartile range, and dots the outliers (> 1.5 times the interquartile range). See Table C3 for statistical values.

**Figure B4.**
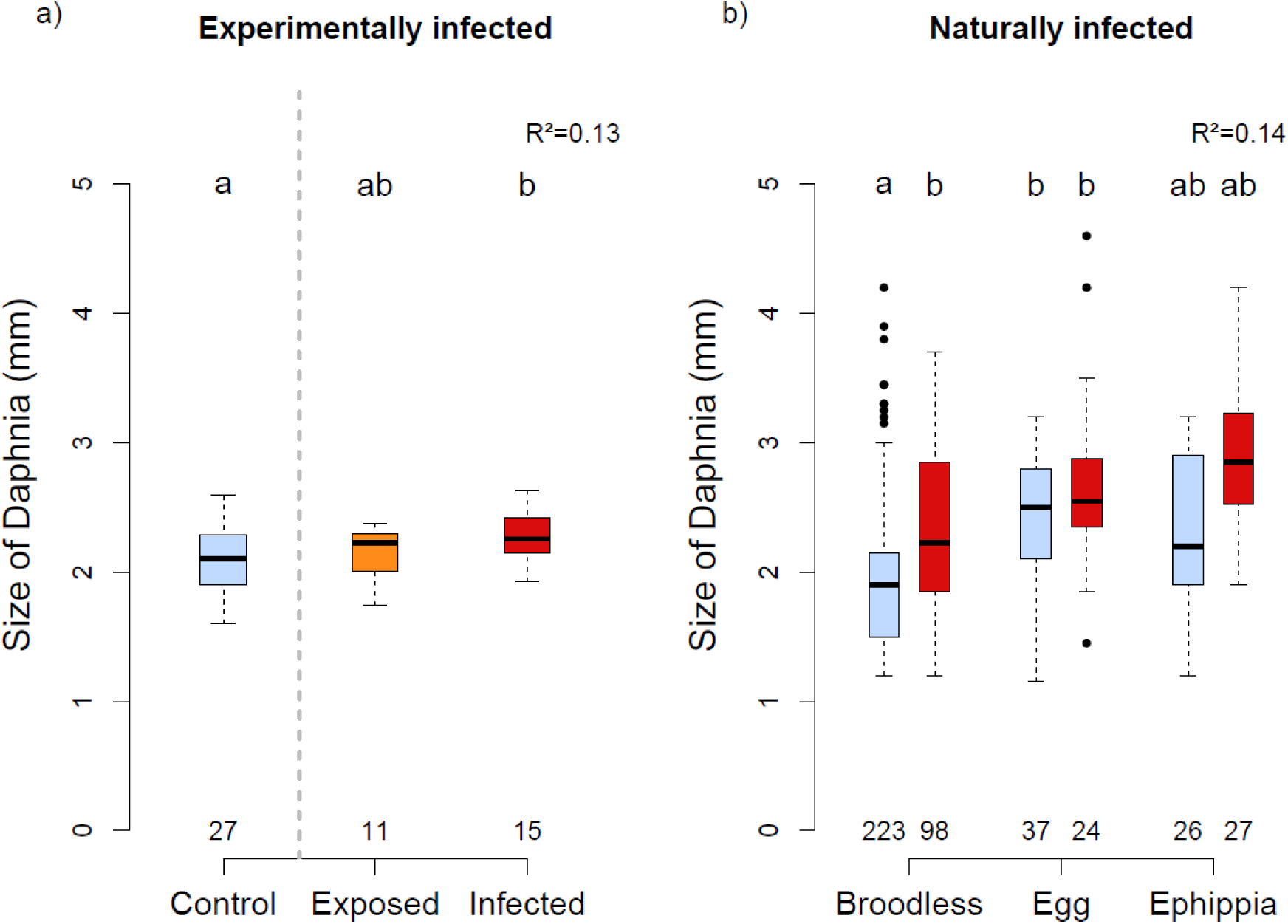
Effects of DIV-1 on host size on a) experimentally infected (healthy/control, exposed, infected); and b) naturally infected *D. magna* (broodless, with eggs, or with ephippia). Numbers are the numbers of *D. magna* for each category. The same letters indicate groups that are not significantly different at 0.05. a) Central bars represent the median, boxes the interquartile range, and dots the outliers (> 1.5 times the interquartile range); and b) dots represent the means and bars the 95% confidence intervals. See Table C4 for statistical values.

## Appendix C: Supplementary figures and tables of statistics

**Figure C1.**
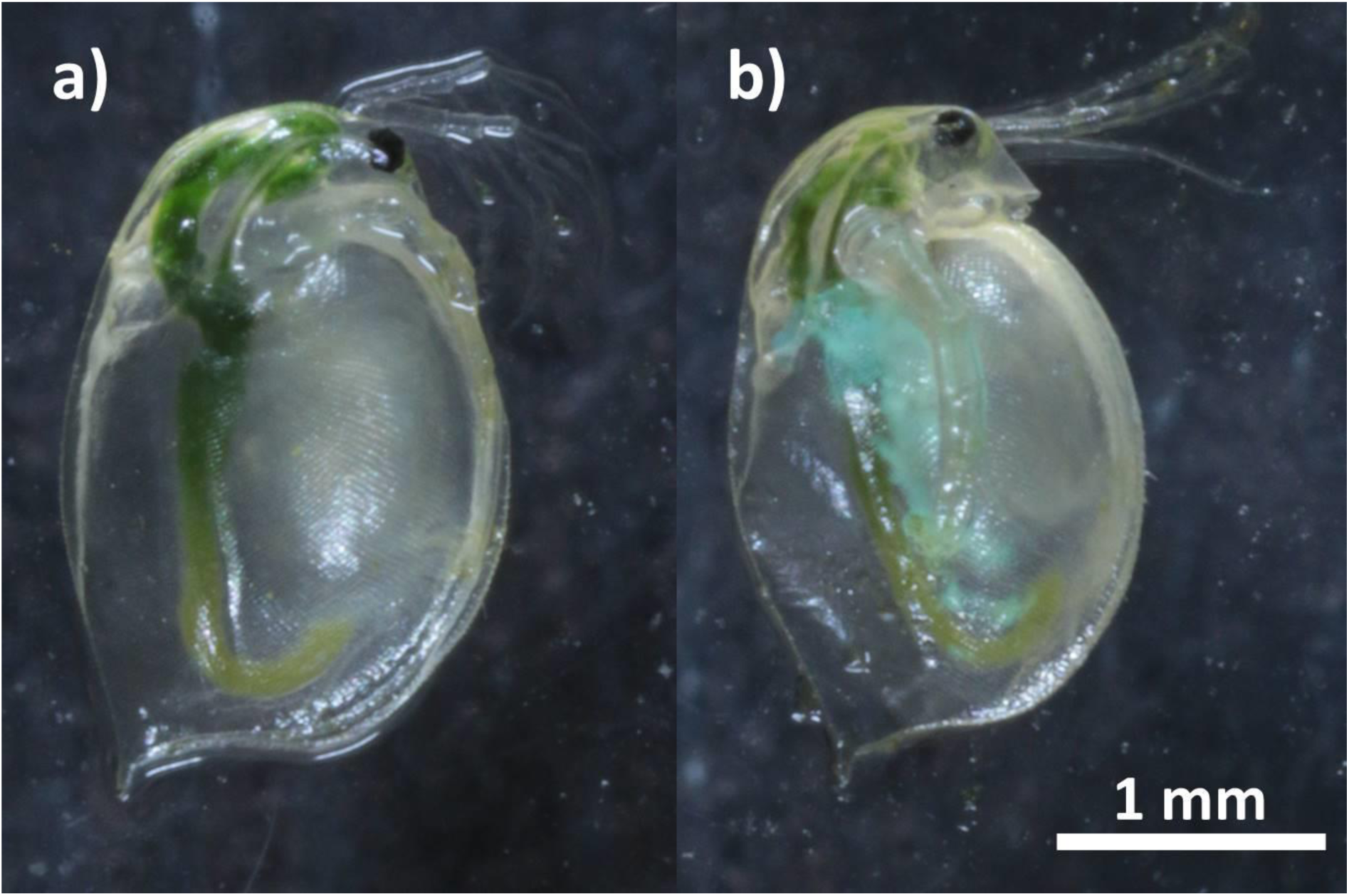
Photographies of *Daphnia magna* from Paris’ pound. a) non-white *D. magna*, b) white *D. magna*. Look the blue-green coloration around the gut (from Prosnier, 2018)

**Figure C2.**
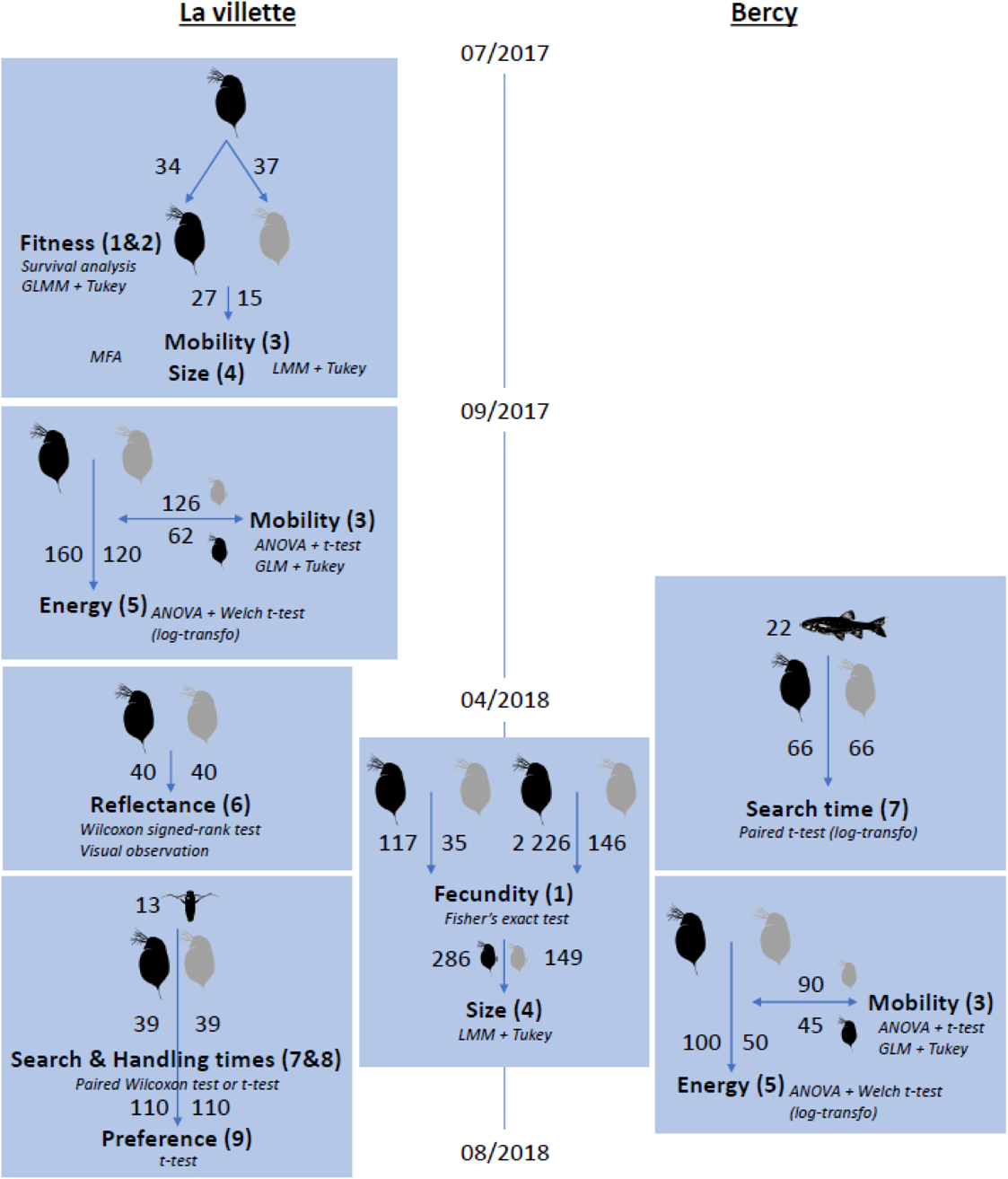
Graphical representation of all measurments performed in the two ponds (La Villette and Bercy) in Paris (France), from July 2017 to August 2018. The measurments are in bold (with their number id in brackets). The statistical tests used are in italic. Numbers are the number of used *D. magna* (non-white daphnia in black and white daphnia in grey), and for predation experiments, the number of fish and *Notonecta*. See also Table 1.

**Figure C3.**
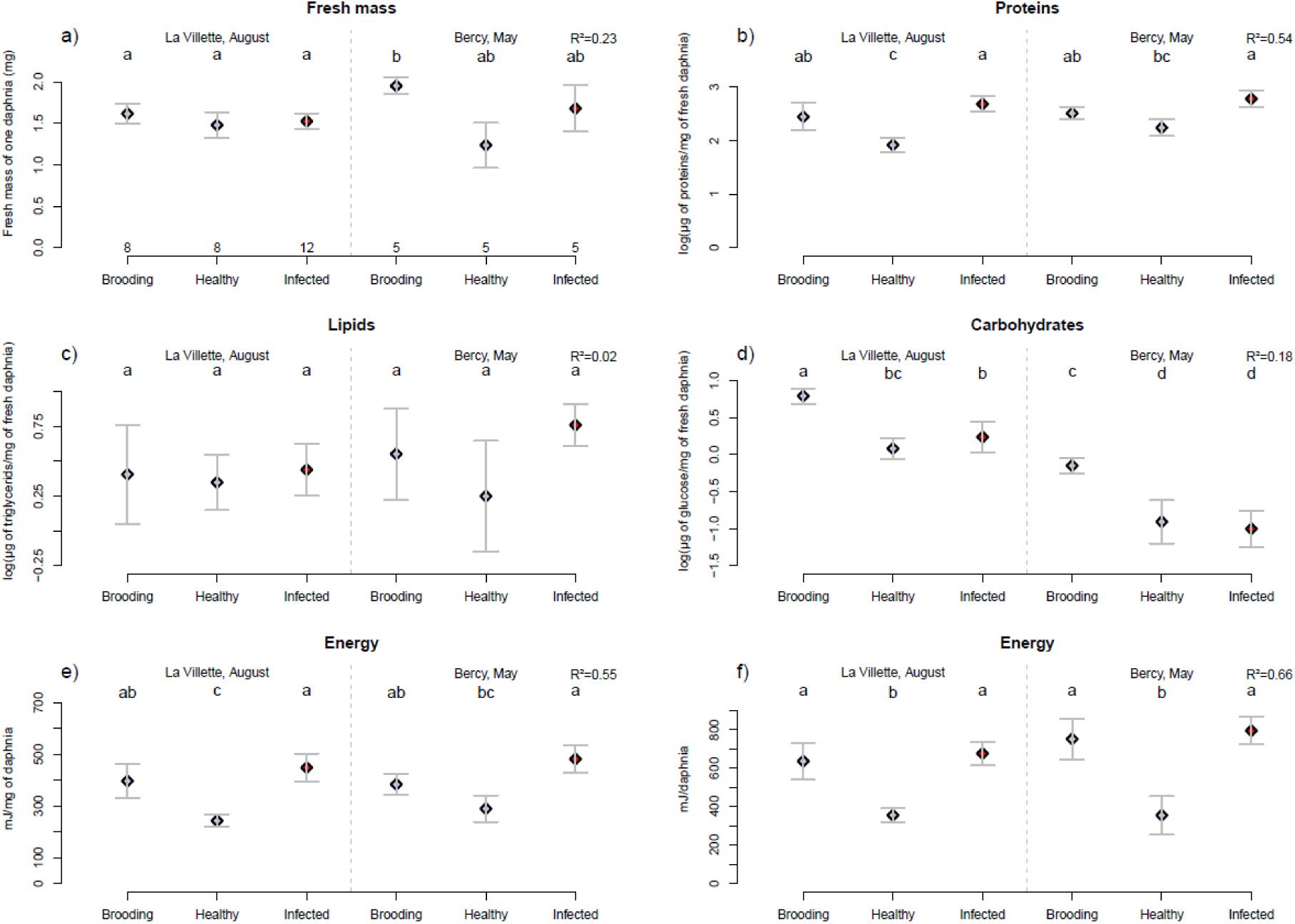
Energy content of *D. magna* for the two populations. a) Biomass, b) protein content, c) lipid content, d) carbohydrate content, e) energy (in mJ) by mg of *D. magna*, and f) energy (in mJ) by *D. magna*. Numbers in a) are the numbers in pools of 10 *D. magna* for each category. The same letters indicate groups that are not significantly different at 0.05. Dots represent the means and bars the 95% confidence intervals. See Table C5 for statistical values.

**Table C1.** Statistical results of DIV-1 effects on fecundity and mortality for the experimental infection (Fig. B1)

|  |  | Mortality |  | Reproduction |  | Fitness |  |  |
| --- | --- | --- | --- | --- | --- | --- | --- | --- |
|  |  | Survival | Adult time | Age at maturity | Mean clutch size | Clutch frequency | Number of clutches | Number of offspring |
| Global effect | df | 2 | 2 | 2 | 2 | 2 | 4 | 2 |
| | $\chi^2$ | 58.3 | 61.7 | 4.6 | 0.222 | 1.356 | 99.542 | 137.52 |
|  | p-value | **< 0.001** | **< 0.001** | 0.102 | 0.895 | 0.508 | **< 0.001** | **< 0.001** |
|  | R<sup>2</sup> | NA | NA | NA | 0 | 0 | 0.8 | 0.73 |
| Control-Infected |  | **< 0.001** | **< 0.001** | **0.037** | 0.940 | 0.728 | **< 0.001** | **< 0.001** |
| Control-Exposed |  | **0.01** | **0.011** | 0.252 | 0.956 | 0.987 | **0.001** | **< 0.001** |
| Exposed-Infected |  | **< 0.001** | **< 0.001** | 0.78 | 0.725 | 0.285 | **< 0.001** | **< 0.001** |

**Table C2.** Statistical results of DIV-1 effects on fecundity for naturally infected *D. magna* (Fig. B2)

| Pond | Bercy |  |  |  | La Villette |  |  |
| --- | --- | --- | --- | --- | --- | --- | --- |
| Date | 19/04 | 03/05 | 17/05 | 19/06 | 03/05 | 17/05 | 19/06 |
| p-value | **0.022** | 0.1 | 0.223 | 0.246 | 0.728 | **0.008** | 0.56 |

**Table C3.** Statistical results of DIV-1 effects on host mobility (Fig. B3)

|  |  | Mean speed | Max speed | Inactivity | Number of turnings |
| --- | --- | --- | --- | --- | --- |
| Experimentally infected |  |  |  |  |  |
| Global effect | df | 2 | 2 | 2 | 2 |
| | $\chi^2$ | 6.206 | 6.538 | 6.530 | 11.589 |
|  | p-value | **0.04** | **0.04** | **0.04** | **0.003** |
|  | R<sup>2</sup> | 0.13 | 0.05 | 0.13 | 0.23 |
| Control-Infected |  | 0.526 | 0.532 | 0.678 | 0.064 |
| Control-Exposed |  | **0.034** | **0.030** | **0.029** | **0.003** |
| Exposed-Infected |  | 0.308 | 0.215 | 0.201 | 0.461 |
| Naturally infected |  |  |  |  |  |
| Global effect | df | 319 | 319 | 3-319 | 319 |
|  | F | NA | NA | 42.32 | NA |
|  | p-value (status) | **< 0.001** | **< 0.001** | **< 0.001** | **< 0.001** |
|  | p-value (pond) | **< 0.001** | **0.004** | **< 0.001** | **< 0.001** |
|  | p-value (status x pond) | 0.17 | **0.002** | **< 0.001** | **0.002** |
|  | R<sup>2</sup> (status) | 0.11 | 0.07 | 0.07 | 0.08 |
| La Villette,<br>August | Healthy-Infected | 0.18 | 0.07 | 0.22 | 0.566 |
| Bercy, May | Healthy-Infected | **< 0.001** | **< 0.001** | **< 0.001** | **< 0.001** |

**Table C4.** Statistical results of DIV-1 effects on host size (Fig. B4)

|  |  | Size |
| --- | --- | --- |
| Experimentally infected |  |  |
| Global effect | df | 2 |
| | $\chi^2$ | 6.3561 |
|  | p-value | **0.042** |
|  | R<sup>2</sup> | 0.13 |
| Control-Infected |  | **0.033** |
| Control-Exposed |  | 0.751 |
| Exposed-Infected |  | 0.294 |
| Natural populations |  |  |
| Global effect | p-value (status) | **<0.001** |
|  | p-value (egg) | **<0.001** |
|  | p-value (status x egg) | 0.902 |
|  | R<sup>2</sup> (status) | 0.14 |
| Healthy-Infected | Broodless | **0.01** |
|  | Egg | 0.22 |
|  | Ephippia | 0.98 |
| Fish predation |  |  |
| Global effect | df | 1 |
| | $\chi^2$ | 0.062296 |
|  | p-value | 0.803 |
|  | R<sup>2</sup> | NA |
| Notonecta predation |  |  |
| Global effect | df | 1-55 |
|  | F-value | 25.49 |
|  | p-value | **<0.001** |
|  | R<sup>2</sup> | 0.32 |

**Table C5.**
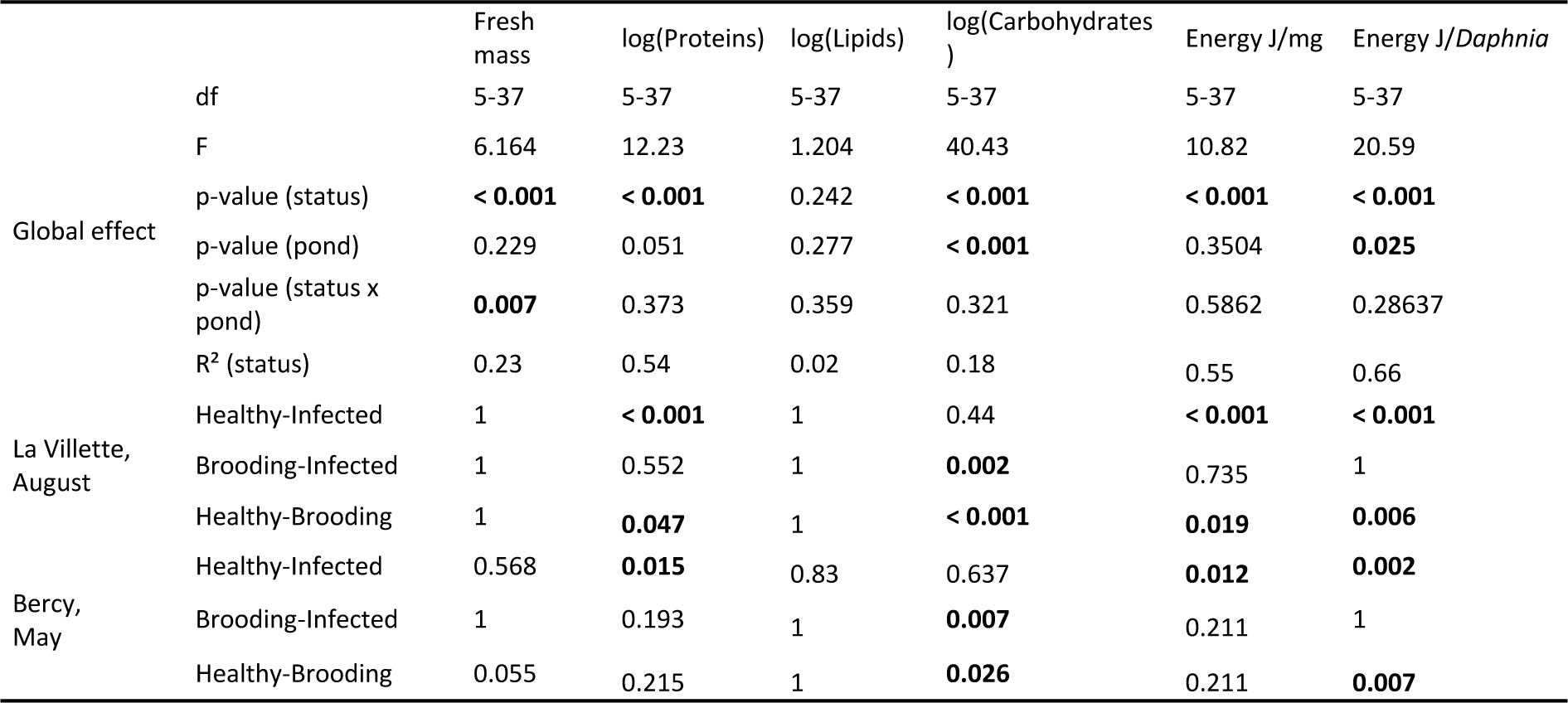
Statistical results of DIV-1 effects on host composition (Fig. C3, Table 2)

**Table C6.** Statistical results of DIV-1 effects on host reflectance (Fig. 2)

|  | UV peak | Blue peak | Orange peak |
| --- | --- | --- | --- |
| df | NA | NA | NA |
| w | 619.5 | 316.5 | 1394 |
| p-value | 0.083 | **< 0.001** | **< 0.001** |

**Table C7.** Statistical results of DIV-1 effects on host vulnerability to predation (Fig. 3 and A1)

|  |  | Search time |  | Handling time | Preference |
| --- | --- | --- | --- | --- | --- |
|  |  | Fish | Notonecta |  |  |
| All catches | df | 53 | NA | NA | 10 |
|  | t/v | 0.58677 | 127 | 6 | 2.1137 |
|  | p-value | 0.28 | 0.891 | **< 0.001** | **0.03** |
| 1st catch | df | 21 | NA | 12 |  |
|  | t/v | 1.8357 | 27 | -4.312 |  |
|  | p-value | **0.04** | 0.9 | **< 0.001** |  |
| 2nd catch | df | 21 | NA | -3.2928 |  |
|  | t/v | -0.77946 | 22 | 8 |  |
|  | p-value | 0.778 | 0.545 | **0.005** |  |
| 3rd catch | df | 9 | NA | 3 |  |
|  | t/v | 0.58129 | 4 | -3.6364 |  |
|  | p-value | 0.288 | 0.687 | **0.018** |  |

**Table C8.** Statistical results of profitability analyses

|  | Profitability vs null model | Null model vs Uniform distribution |
| --- | --- | --- |
| D | 0.9054 | 0.0132 |
| p-value | **< 0.001** | 0.347 |

## References

1. Agnew P, Bedhomme S, Haussy C, Michalakis Y (1999) Age and size at maturity of the mosquito Culex pipiens infected by the microsporidian parasite Vavraia culicis. Proceedings of the Royal Society of London. Series B: Biological Sciences, 266, 947–952. https://doi.org/10.1098/rspb.1999.0728

2. Amoros C (1984) Introduction pratique à la systématique des organismes des eaux continentales françaises. Bulletin mensuel de la Société Linnéenne de Lyon, 53, 72–145.

3. Bedhomme S, Agnew P, Sidobre C, Michalakis Y (2004) Virulence reaction norms across a food gradient. Proceedings of the Royal Society B: Biological Sciences, 271, 739–744. https://doi.org/10.1098/rspb.2003.2657

4. Bennett RR, Ruck P (1970) Spectral sensitivities of dark- and light-adapted Notonecta compound eyes. Journal of Insect Physiology, 16, 83–88. https://doi.org/10.1016/0022-1910(70)90115-0

5. Bethel WM, Holmes JC (1977) Increased vulnerability of amphipods to predation owing to altered behavior induced by larval acanthocephalans. Canadian journal of zoology, 55, 110–115. https://doi.org/10.1139/z77-013

6. Bland M (2013) Do baseline p-values follow a uniform distribution in randomised trials? (M Law, Ed,). PLoS ONE, 8, e76010. https://doi.org/10.1371/journal.pone.0076010

7. Bownik A (2017) Daphnia swimming behaviour as a biomarker in toxicity assessment: A review. Science of The Total Environment, 601–602, 194–205. https://doi.org/10.1016/j.scitotenv.2017.05.199

8. Bradford MM (1976) A rapid and sensitive method for the quantitation of microgram quantities of protein utilizing the principle of protein-dye binding. Analytical Biochemistry, 72, 248–254. https://doi.org/10.1006/abio.1976.9999

9. Britton NF, Jane White KA (2021) The effect of covert and overt infections on disease dynamics in honey-bee colonies. Bulletin of Mathematical Biology, 83, 67. https://doi.org/10.1007/s11538-021-00892-6

10. Cáceres CE, Knight CJ, Hall SR (2009) Predator-spreaders: Predation can enhance parasite success in a planktonic host-parasite system. Ecology, 90, 2850–2858. https://doi.org/10.1890/08-2154.1

11. Cézilly F, Favrat A, Perrot-Minnot M-J (2013) Multidimensionality in parasite-induced phenotypic alterations: ultimate versus proximate aspects. Journal of Experimental Biology, 216, 27–35. https://doi.org/10.1242/jeb.074005

12. Chadwick W, Little TJ (2005) A parasite-mediated life-history shift in Daphnia magna. Proceedings of the Royal Society B: Biological Sciences, 272, 505–509. https://doi.org/10.1098/rspb.2004.2959

13. Chang K-H, Hanazato T (2003) Vulnerability of cladoceran species to predation by the copepod Mesocyclops leuckarti: laboratory observations on the behavioural interactions between predator and prey. Freshwater Biology, 48, 476–484. https://doi.org/10.1046/j.1365-2427.2003.01021.x

14. Charnov EL (1976a) Optimal foraging, the marginal value theorem. Theoretical population biology, 9, 129–136. https://doi.org/10.1016/0040-5809(76)90040-X

15. Charnov EL (1976b) Optimal foraging: Attack strategy of a mantid. The American Naturalist, 110, 141– 151.

16. de Coen WM, Janssen CR (1997) The use of biomarkers in Daphnia magna toxicity testing. IV. Cellular Energy Allocation: a new methodology to assess the energy budget of toxicant-stressed Daphnia populations. Journal of Aquatic Ecosystem Stress and Recovery, 6, 43–55.

17. Cumming G, Finch S (2005) Inference by Eye: Confidence Intervals and How to Read Pictures of Data. American Psychologist, 60, 170–180. https://doi.org/10.1037/0003-066X.60.2.170

18. Dallas T, Holtackers M, Drake JM (2016) Costs of resistance and infection by a generalist pathogen. Ecology and Evolution, 6, 1737–1744. https://doi.org/10.1002/ece3.1889

19. Decaestecker E, Declerck SAJ, De Meester L, Ebert D (2005) Ecological implications of parasites in natural *Daphnia* populations. Oecologia, 144, 382–390. https://doi.org/10.1007/s00442-005-0083-7

20. Decaestecker E, Vergote A, Ebert D, De Meester L (2003) Evidence for strong host clone-parasite species interactions in the Daphnia microparasite system. Evolution, 57, 784–792. https://doi.org/10.1554/0014-3820(2003)057[0784:EFSHCS]2.0.CO;2

21. Decaestecker E, Verreydt D, De Meester L, Declerck SAJ (2015) Parasite and nutrient enrichment effects on Daphnia interspecific competition. Ecology, 96, 1421–1430. https://doi.org/10.1890/14-1167.1

22. Dobson AP, Lafferty KD, Kuris AM, Hechinger RF, Jetz W (2008) Homage to Linnaeus: How many parasites? How many hosts? Proceedings of the National Academy of Sciences, 105, 11482–11489. https://doi.org/10.1073/pnas.0803232105

23. Dodson S, Ramcharan C (1991) Size-specific swimming behavior of Daphnia pulex. Journal of Plankton Research, 13, 1367–1379. https://doi.org/10.1093/plankt/13.6.1367

24. Ebert D (2005) *Ecology, epidemiology and evolution of parasitism in* Daphnia. Bethesda (MD).

25. Ebert D (2022) Daphnia as a versatile model system in ecology and evolution. EvoDevo, 13, 16. https://doi.org/10.1186/s13227-022-00199-0

26. Ebert D, Lipsitch M, Mangin KL (2000) The effect of parasites on host population density and extinction: Experimental epidemiology with Daphnia and six microparasites. The American Naturalist, 156, 459–477. https://doi.org/10.1086/303404

27. Emlen JM (1966) The role of time and energy in food preference. The American Naturalist, 100, 611– 617. https://doi.org/10.1086/282455

28. Flick AJ, Acevedo MA, Elderd BD (2016) The negative effects of pathogen-infected prey on predators: a meta-analysis. Oikos, 125, 1554–1560. https://doi.org/10.1111/oik.03458

29. Foray V, Pelisson PF, Bel-Venner MC, Desouhant E, Venner S, Menu F, Giron D, Rey B (2012) A handbook for uncovering the complete energetic budget in insects: the van Handel’s method (1985) revisited. Physiological Entomology, 37, 295–302. https://doi.org/10.1111/j.1365-3032.2012.00831.x

30. Forshay KJ, Johnson PTJ, Stock M, Peñalva C, Dodson SI (2008) Festering food: chytridiomycete pathogen reduces quality of Daphnia host as a food resource. Ecology, 89, 2692–2699. https://doi.org/10.1890/07-1984.1

31. Frainer A, McKie BG, Amundsen P-A, Knudsen R, Lafferty KD (2018) Parasitism and the biodiversity-functioning relationship. Trends in Ecology & Evolution, 33, 260–268. https://doi.org/10.1016/j.tree.2018.01.011

32. Gerritsen J, Strickler JR (1977) Encounter probabilities and community structure in zooplancton: a mathematical model. Journal of the Fisheries Research Board of Canada, 34, 73–82.

33. Giller PS (1986) The natural diet of the Notonectidae: field trials using electrophoresis. Ecological Entomology, 11, 163–172. https://doi.org/10.1111/j.1365-2311.1986.tb00291.x

34. Gnaiger E (1983) Calculation of energetic and biochemical equivalents. In: Polarographic Oxygen Sensors. Aquatic and Physiological Applications. (eds Gnaiger E, Forstner H), pp. 337–345. Springer Verlag, Berlin.

35. Goren L, Ben-Ami F (2017) To eat or not to eat infected food: a bug’s dilemma. Hydrobiologia, 798, 25–32. https://doi.org/10.1007/s10750-015-2373-3

36. Green J (1974) Parasites and epibionts of Cladocera. The Transactions of the Zoological Society of London, 32, 417–515. https://doi.org/10.1111/j.1096-3642.1974.tb00031.x

37. Hall SR, Becker CR, Cáceres CE (2007) Parasitic castration: a perspective from a model of dynamic energy budgets. Integrative and Comparative Biology, 47, 295–309. https://doi.org/10.1093/icb/icm057

38. Hudson PJ, Dobson AP, Newborn D (1992) Do parasites make prey vulnerable to predation? Red grouse and parasites. The Journal of Animal Ecology, 61, 681. https://doi.org/10.2307/5623

39. Hülsmann S, Weiler W (2000) Adult, not juvenile mortality as a major reason for the midsummer decline of a Daphnia population. Journal of Plankton Research, 22, 151–168. https://doi.org/10.1093/plankt/22.1.151

40. Jacquin L, Mori Q, Médoc V (2013) Does the carotenoid-based colouration of Polymorphus minutus facilitate its trophic transmission to definitive hosts? Parasitology, 140, 1310–1315. https://doi.org/10.1017/S0031182013000760

41. Jacquin L, Mori Q, Pause M, Steffen M, Médoc V (2014) Non-specific manipulation of gammarid behaviour by P. minutus parasite enhances their predation by definitive bird hosts (CG de Leaniz, Ed,). PLoS ONE, 9, e101684. https://doi.org/10.1371/journal.pone.0101684

42. Johnson PTJ, Stanton DE, Preu ER, Forshay KJ, Carpenter SR (2006) Dining on disease: how interactions between infection and environment affect predation risk. Ecology, 87, 1973–80. https://doi.org/10.1890/0012-9658(2006)87[1973:DODHIB]2.0.CO;2

43. Kondoh M (2003) Foraging adaptation and the relationship between food-web complexity and stability. Science, 299, 1388–1391. https://doi.org/10.1126/science.1079154

44. Labbé P, Vale PF, Little TJ (2010) Successfully resisting a pathogen is rarely costly in Daphnia magna. BMC Evolutionary Biology, 10, 355. https://doi.org/10.1186/1471-2148-10-355

45. Lampert W, Sommer U (2007) Limnoecology. Oxford Biology.

46. Van der Lee GH, Vonk JA, Verdonschot RCM, Kraak MHS, Verdonschot PFM, Huisman J (2021) Eutrophication induces shifts in the trophic position of invertebrates in aquatic food webs. Ecology, 102, 1–13. https://doi.org/10.1002/ecy.3275

47. Lefèvre T, Lebarbenchon C, Gauthier-Clerc M, Missé D, Poulin R, Thomas F (2009) The ecological significance of manipulative parasites. Trends in Ecology & Evolution, 24, 41–48. https://doi.org/10.1016/j.tree.2008.08.007

48. MacArthur RH, Pianka ER (1966) On optimal use of a patchy environment. The American Naturalist, 100, 603–609. https://doi.org/10.1086/282454

49. Manly BFJ (1974) A Model for Certain Types of Selection Experiments. Biometrics, 30, 281–294. https://doi.org/10.2307/2529649

50. Marina CF, Arredondo-Jiménez JI, Castillo A, Williams T (1999) Sublethal effects of iridovirus disease in a mosquito. Oecologia, 119, 383–388. https://doi.org/10.1007/s004420050799

51. Marina CF, Ibarra JE, Arredondo-Jimenez JI, Fernandez-Salas I, Valle J, Williams T (2003) Sublethal iridovirus disease of the mosquito Aedes aegypti is due to viral replication not cytotoxicity. Medical and Veterinary Entomology, 17, 187–194. https://doi.org/10.1046/j.1365-2915.2003.00422.x

52. McCann KS (2000) The diversity-stability debate. Nature, 405, 228–233. https://doi.org/10.1038/35012234

53. McTaggart SJ, Conlon C, Colbourne JK, Blaxter ML, Little TJ (2009) The components of the Daphnia pulex immune system as revealed by complete genome sequencing. BMC Genomics, 10, 175. https://doi.org/10.1186/1471-2164-10-175

54. Médoc V, Piscart C, Maazouzi C, Simon L, Beisel J-N (2011) Parasite-induced changes in the diet of a freshwater amphipod: field and laboratory evidence. Parasitology, 138, 537–546. https://doi.org/10.1017/S0031182010001617

55. Modarressie R, Rick IP, Bakker TCM (2013) Ultraviolet reflection enhances the risk of predation in a vertebrate. Current Zoology, 59, 151–159. https://doi.org/10.1093/czoolo/59.2.151

56. Newey S, Thirgood S (2004) Parasite–mediated reduction in fecundity of mountain hares. Proceedings of the Royal Society of London. Series B: Biological Sciences, 271, S413–S415. https://doi.org/10.1098/rsbl.2004.0202

57. O’Keefe TC, Brewer MC, Dodson SI (1998) Swimming behavior of Daphnia : its role in determining predation risk. Journal of Plankton Research, 20, 973–984. https://doi.org/10.1093/plankt/20.5.973

58. Otti O, Gantenbein-Ritter I, Jacot A, Brinkhof MWG (2012) Immune response increases predation risk. Evolution, 66, 732–739. https://doi.org/10.1111/j.1558-5646.2011.01506.x

59. Ouisse T, Laparie M, Lebouvier M, Renault D (2017) New insights into the ecology of Merizodus soledadinus, a predatory carabid beetle invading the sub-Antarctic Kerguelen Islands. Polar Biology, 40, 2201–2209. https://doi.org/10.1007/s00300-017-2134-z

60. Packer C, Holt RD, Hudson PJ, Lafferty KD, Dobson AP (2003) Keeping the herds healthy and alert: implications of predator control for infectious disease. Ecology Letters, 6, 797–802. https://doi.org/10.1046/j.1461-0248.2003.00500.x

61. Peterson RO, Page RE (1988) The rise and fall of isle royale wolves, 1975-1986. Journal of Mammalogy, 69, 89–99. https://doi.org/10.2307/1381751

62. Plaistow SJ, Troussard J-P, Cézilly F (2001) The effect of the acanthocephalan parasite Pomphorhynchus laevis on the lipid and glycogen content of its intermediate host Gammarus pulex. International Journal for Parasitology, 31, 346–351. https://doi.org/10.1016/S0020-7519(01)00115-1

63. Prins HHT, Weyerhaeuser FJ (1987) Epidemics in populations of wild ruminants: Anthrax and impala, rinderpest and buffalo in Lake Manyara National Park, Tanzania. Oikos, 49, 28–38. https://doi.org/10.2307/3565551

64. Prosnier L (2018) Implications écologiques et évolutives du parasitisme sur les structures trophiques. Sorbonne Université.

65. Prosnier L, Loeuille N, Hulot FD, Renault D, Piscart C, Bicocchi B, Deparis M, Lam M, Médoc V (2022) Datasets and R source code of manuscript “Parasites make hosts more profitable but less available to predators.” Dataset on Zenodo. https://doi.org/10.5281/zenodo.6006617

66. Prosnier L, Médoc V, Loeuille N (2018) Parasitism effects on coexistence and stability within simple trophic modules. Journal of Theoretical Biology, 458, 68–77. https://doi.org/10.1016/j.jtbi.2018.09.004

67. Prosnier L, Médoc V, Loeuille N (2020) Evolution of predator foraging in response to prey infection favors species coexistence. *bioRxiv*, 2020.04.18.047811. https://doi.org/10.1101/2020.04.18.047811

68. Read AF (1994) The evolution of virulence. Trends in Microbiology, 2, 73–76. https://doi.org/10.1016/0966-842X(94)90537-1

69. Reynolds CS (2011) Daphnia: Development of Model Organism in Ecology and Evolution -2011 Winfried Lampert (2011) Excellence in Ecology Series. Freshwater Reviews, 4, 85–87. https://doi.org/10.1608/FRJ-4.1.425

70. Riessen HP (1999) Predator-induced life history shifts in Daphnia : a synthesis of studies using meta-analysis. Canadian Journal of Fisheries and Aquatic Sciences, 56, 2487–2494. https://doi.org/10.1139/f99-155

71. Rosa RD, Barracco MA (2010) Antimicrobial peptides in crustaceans. Invertebrate Survival Journal, 7, 262–284.

72. Sánchez CA, Becker DJ, Teitelbaum CS, Barriga P, Brown LM, Majewska AA, Hall RJ, Altizer S (2018) On the relationship between body condition and parasite infection in wildlife: a review and meta-analysis (J Davies, Ed,). Ecology Letters, 21, 1869–1884. https://doi.org/10.1111/ele.13160

73. Schwartz SS, Cameron GN (1993) How do parasites cost their hosts? Preliminary answers from trematodes and Daphnia obtusa. Limnology and Oceanography, 38, 602–612. https://doi.org/10.4319/lo.1993.38.3.0602

74. Sorrell I, White A, Pedersen AB, Hails RS, Boots M (2009) The evolution of covert, silent infection as a parasite strategy. Proceedings of the Royal Society B: Biological Sciences, 276, 2217–2226. https://doi.org/10.1098/rspb.2008.1915

75. Spitze K (1985) Functional response of an ambush predator: Chaoborus americanus predation on Daphnia pulex. Ecology, 66, 938–949. https://doi.org/10.2307/1940556

76. Stibor H, Lampert W (1993) Estimating the size at maturity in field populations of Daphnia (Cladocera). Freshwater Biology, 30, 433–438. https://doi.org/10.1111/j.1365-2427.1993.tb00826.x

77. Thomas F, Poulin R, Brodeur J (2010) Host manipulation by parasites: a multidimensional phenomenon. Oikos, 119, 1217–1223. https://doi.org/10.1111/j.1600-0706.2009.18077.x

78. Toenshoff ER, Fields PD, Bourgeois YX, Ebert D (2018) The end of a 60-year riddle: Identification and genomic characterization of an iridovirus, the causative agent of white fat cell disease in zooplankton. G3 Genes|Genomes|Genetics, 8, 1259–1272. https://doi.org/10.1534/g3.117.300429

79. Untersteiner H, Kahapka J, Kaiser H (2003) Behavioural response of the cladoceran Daphnia magna STRAUS to sublethal Copper stress - Validation by image analysis. Aquatic Toxicology, 65, 435–442. https://doi.org/10.1016/S0166-445X(03)00157-7

80. Vance SA, Peckarsky BL (1997) The effect of mermithid parasitism on predation of nymphal Baetis bicaudatus (Ephemeroptera) by invertebrates. Oecologia, 110, 147–152. https://doi.org/10.1007/s004420050143

81. Williams T (1993) Covert iridovirus infection of blackfly larvae. Proceedings of the Royal Society of London. Series B: Biological Sciences, 251, 225–230. https://doi.org/10.1098/rspb.1993.0033

82. Williams T (2008) Natural invertebrate hosts of iridoviruses (Iridoviridae). Neotropical Entomology, 37, 615–632. https://doi.org/10.1590/S1519-566X2008000600001

83. Williams T, Barbosa-Solomieu V, Chinchar VG (2005) A decade of advances in iridovirus research. Advances in Virus Research, 65, 173–248. https://doi.org/10.1016/S0065-3527(05)65006-3

84. Xie H, Wei J, Qin Q (2016) Antiviral function of Tachyplesin I against iridovirus and nodavirus. Fish & Shellfish Immunology, 58, 96–102. https://doi.org/10.1016/j.fsi.2016.09.015

